# Australian terrestrial environments harbour extensive RNA virus diversity

**DOI:** 10.1101/2023.10.09.561620

**Authors:** Sabrina Sadiq, Erin Harvey, Jonathon C. O. Mifsud, Budiman Minasny, Alex. B. McBratney, Liana E. Pozza, Jackie E. Mahar, Edward C. Holmes

## Abstract

Australia is home to a diverse range of unique native fauna and flora. To address whether Australian ecosystems also harbour unique viruses, we performed meta-transcriptomic sequencing of 16 farmland and sediment samples taken from the east and west coasts of Australia. We identified 2,562 putatively novel viruses across 15 orders, the vast majority of which belonged to the microbe-associated phylum *Lenarviricota*. In many orders, the novel viruses identified here comprised entirely new clades, such as the *Nodamuvirales* and *Ghabrivirales*. Novel viruses also fell between established genera or families, such as in the *Cystoviridae* and *Picornavirales*, while highly divergent lineages were identified in the *Martellivirales* and *Ghabrivirales*. Viral abundance and alpha diversity were influenced by sampling site, soil type and land use, but not by depth from the surface. In sum, Australian soils and sediments are home to remarkable viral diversity, reflecting the biodiversity of local fauna and flora.

## INTRODUCTION

RNA viruses are ubiquitous, diverse, and can play important roles in multiple ecological processes. Yet a strong focus on viruses of clinical or agricultural significance has limited our understanding of RNA viruses as potentially key components of global ecosystems (French and Holmes, 2020). Similarly, studies of environmental viruses have largely considered marine systems and typically characterised DNA viromes (Trubl *et al*., 2020), leaving the terrestrial RNA virome understudied (Starr *et al*., 2019; Chen *et al*., 2022; Hillary *et al*., 2022).

Soil environments are complex and diverse, host to an intricate network of macro- and micro-organisms that form unique ecosystems with crucial global functions. An estimated 10^10^ microbes are present in one gram of soil, with species diversity up to the tens of thousands (Raynaud and Nunan, 2014). Microbial community compositions both affect and depend on factors such as the physicochemical properties of soil (Singh *et al*., 2009; Dequiedt *et al*., 2011), land use (Tian *et al*., 2017), and depth (Xue *et al*., 2022). The abundance and diversity of organisms in soil implies that there must also be abundant and diverse viruses infecting these hosts, which may play their own roles in soil cycles and maintenance. Indeed, viruses in soil have been estimated from below detection limits in hot deserts to over 10^10^/g soil in wetlands (Williamson et al., 2017). Viruses have documented roles in carbon metabolism (Trubl *et al*., 2018; Jin *et al*., 2019; Starr *et al*., 2019) and phosphorus metabolism (Han *et al*., 2022), as well as gene transfer to their bacterial hosts, aiding in host extremotolerance and adaptation to environmental stressors (Hwang *et al*., 2021; Huang *et al*., 2021).

Recent metatranscriptomic (i.e., total RNA-sequencing) studies have led to the discovery and characterisation of novel viruses from diverse ecosystems, including aquatic environments (Wolf *et al*., 2020) and soil (Chen *et al*., 2022; Hillary *et al*., 2022). In particular, metatranscriptomic analyses of diverse soils and freshwater sediments have shown that non-marine environments are a rich source of viral diversity, with thousands of novel RNA viruses identified in every major lineage of RNA viruses (Chen *et al*., 2022; Hillary *et al*., 2022). Hence, soils provide a valuable means to characterise more of the terrestrial RNA virome. These studies have also led to a deeper understanding of RNA viruses in a broader ecological context, revealing the impacts of human land use, physicochemical properties, and geographical features of the sampling environment on viral abundance and diversity (French *et al*., 2022; Chen *et al*., 2022; Hillary *et al*., 2022).

Australia has been isolated from other continents for tens of millions of years and, as such, has developed many diverse and unique biomes. Flora and fauna have adapted to the continent’s flat, dry, fire-prone, and nutrient-poor landmass, resulting in a remarkable level of biodiversity that is unique to Australia (Steffen, 2009). For example, many native Australian plants have hardened foliage (sclerophylly) and evergreen characteristics, causing herbivores to have slower metabolisms and reptiles to have predominantly invertebrate-based diets (Steffen, 2009). Insects play an active role in the dispersal of seeds and are key consumers of leaves in the absence of any native ruminant species (Steffen, 2009). The majority of Australia’s mammals (87%), reptiles (93%), frogs (94%), and vascular plants (92%) found across the country are endemic (Chapman, 2009). Australia is also a “megadiverse” country, one of 17 that together comprise over 70% of the world’s total biodiversity (Williams *et al*., 2001). With so much diversity in its animals, plants, and environments, it can be assumed that Australian microbes - including viruses - harbour similar levels of diversity.

Little is known about the RNA viromes of Australian soil and sediment systems. Herein, we aimed to provide an initial snapshot of the diversity, abundance, and composition of RNA viruses in Australian environmental samples. In particular, we asked whether the unique flora and fauna of Australia is reflected in a unique soil virome. Accordingly, we performed meta-transcriptomic sequencing of 16 geographically and ecologically distinct farmland soil and riverbank sediment samples taken from New South Wales (NSW) and Western Australia (WA), respectively, two Australian states separated by approximately 3,000 kilometres.

## 2 MATERIALS AND METHODS

### 2.1 Sample collection

Soil samples from NSW were collected in December 2021 from vertosol, sodosol, and chromosol soil types (Isbell, 2016) in cropping/pasture fields and native vegetation at depths of 0-5 cm, 5-15 cm, and 15-30 cm at each site. NSW samples were collected from the University of Sydney-owned ‘Nowley Farm’ on the Liverpool plains due to the presence of multiple soil types and land uses across the farm. In the case of WA, sediment samples were collected in triplicate during April and May 2022 from four points along the riverbanks of the Swan River, Canning River, and Denmark River. Maps of the NSW and WA sampling sites are shown in Fig. 1, with more detail on the properties of the sampling sites provided in Table S1. All samples were packed into sterile 50 ml conical tubes and either stored on ice (NSW samples) or between −30°C and −15°C (WA samples) until transported to a −80°C freezer for long-term storage.

**Figure 1.**
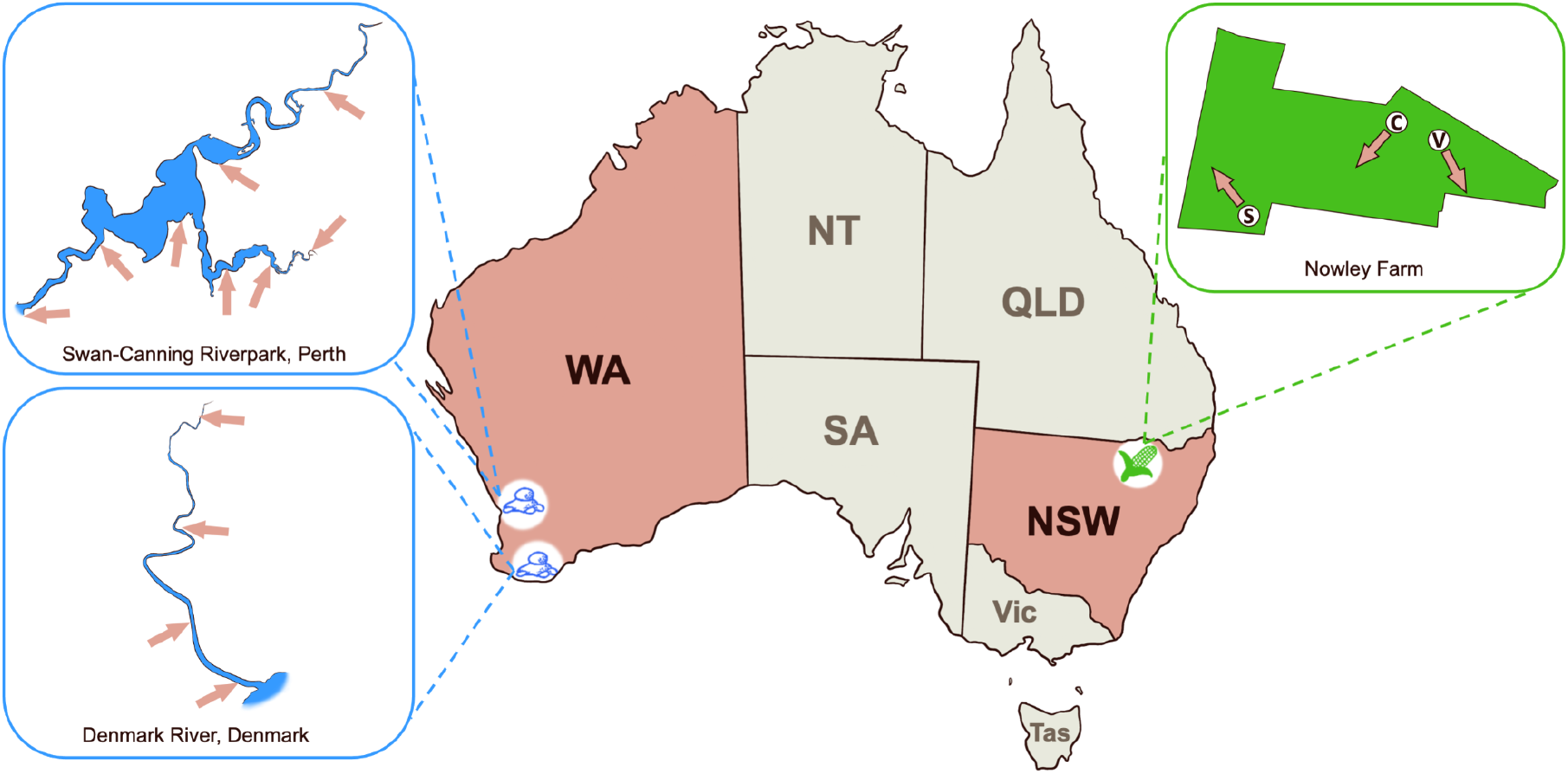
Map of Australia showing riverbank sediment sampling sites along the Swan-Canning Riverpark (Perth, WA) and Denmark River (Denmark, WA), as well as farmland soil sampling sites in Nowley Farm, NSW. In the inset boxes, each sampling site is indicated by an arrow. In the case of the Nowley Farm samples, soil type is indicated by a letter: V - vertosol, C - chromosol, S - sodosol.

### 2.2 RNA extraction, library construction, and sequencing

Total RNA was extracted from the samples using the RNeasy PowerSoil Total RNA Kit (Qiagen) as per the manufacturer’s instructions. Extracted RNA was quantified using the Qubit RNA high sensitivity (HS) Assay Kit on the Qubit Fluorometer v3.0 (Thermo Fisher Scientific) and stored at −80°C prior to library construction and sequencing. Libraries were constructed using the Illumina Stranded Total RNA library preparation protocol and rRNA was removed using the Ribo-Zero Plus rRNA depletion kit (Illumina). Libraries were sequenced on the Illumina NovaSeq 6000 platform (paired-end, 150 bp). Library preparation and sequencing was performed by the Australian Genome Research Facility (AGRF).

### 2.3 Data processing and abundance measurements

Sequence reads were adaptor- and quality-trimmed using Trimmomatic (v0.38) (Bolger *et al*., 2014), then assembled into contigs using MEGAHIT (v1.2.9) (Li *et al*., 2015) employing default assembly parameters. No eukaryotic or bacterial reads were filtered prior to assembly. Sequence quality was checked for both raw reads and trimmed reads using FastQC (v0.11.8) (Andrews, 2010). Assembled contigs were then compared to an in-house curated database of viral sequences from NCBI’s protein sequence database (created in 2019, updated in 2022) using DIAMOND BLASTX (v2.0.9) (Buchfink *et al*., 2015) with an e-value of 1×10^−5^ for sensitivity. Contigs returning positive hits were compared to the non-redundant (nr) protein database as of June 2022, using DIAMOND BLASTX to identify false positives. Contigs with hits to viral sequences were retained and sorted by the taxonomy of the closest relative to the level of virus family (or to the most specific available taxonomic level for divergent and unclassified taxa).

Viral abundance within each library was estimated using RSEM (v1.3.1) (Li and Dewey, 2011), calculated as the expected count of viral contigs divided by the total raw read count x 100. Virome compositions for each library were calculated as the expected count of contigs aligning to each viral family/order/phylum as a proportion of the total expected count of contigs aligning to sequences from the *Riboviria* (RNA viruses). Alpha diversity was described by: (i) richness, or the number of viral taxa in each library, (ii) Shannon diversity, which accounts for both richness and the evenness of their distribution, and (iii) “true” diversity or effective Shannon diversity, calculated as the exponential of each respective Shannon diversity index. These indices were calculated in RStudio (v4.1.1717) (RStudio Team, 2019), R (v4.1.0) (R Core Team, 2021) using an adaptation of the Rhea alpha diversity script (Lagkouvardos *et al*., 2017; Wille *et al*., 2019). Abundance and alpha diversity figures were generated in R using ggplot2 (v3.3.3) (Wickham, 2016). Soil characteristics were tested for their influence on viral abundance and alpha diversity using generalised linear models. The significance of these models was evaluated using χ^2^ tests and significant differences between pairs of groups within each ecological property were determined using post-hoc Tukey tests. To determine if soil virus abundance and alpha diversity were shaped by soil type, land use, or both, we conducted a best-subsets regression analysis. Models were evaluated using the Akaike information criterion (AIC) and the model with the lowest AIC was selected as the best-fit model. Soil characteristics included soil type (chromosol versus sodosol), land use (native vegetation, cropping, and pasture), environment (a combination of soil type and land use) and depth from the surface. As sequence data was obtained from only one depth measurement for chromosol and sediment environments, only sodosol libraries were included in the comparison of viral abundance and alpha diversity across different depths. Viral abundance and alpha diversity were compared between the different sampling sites for sediment libraries. Due to the numerous confounding factors that may influence virus abundance and alpha diversity, including the surrounding environment, geographical location, climate, and storage conditions prior to extraction, detailed comparisons between the NSW farmland soil and WA sediment samples were not conducted.

### 2.4 Identification of novel viruses and phylogenetic analysis

Contigs with DIAMOND BLASTX hits to the viral RNA-dependent RNA polymerase (RdRp) that were greater than 600 nucleotides (nt) in length (arbitrarily set so as to minimise inaccurate classification) and with less than 99% amino acid identity to their closest previously published relative were translated to amino acid sequences. The standard genetic code (i.e., code table 1) was used in most cases, with the exception of 126 sequences from the family *Mitoviridae* (phylum *Lenarviricota*) for which the mitochondrial genetic code (i.e., code table 4) is more likely to be biologically accurate, and indeed provided ORFs of expected lengths where the standard code led to truncation. Translated sequences were checked for the presence of the conserved A, B, and C motifs that characterise viral RdRp. Contigs fulfilling these conditions were included for phylogenetic analysis as these likely represent RNA virus sequences.

**Table 1.**
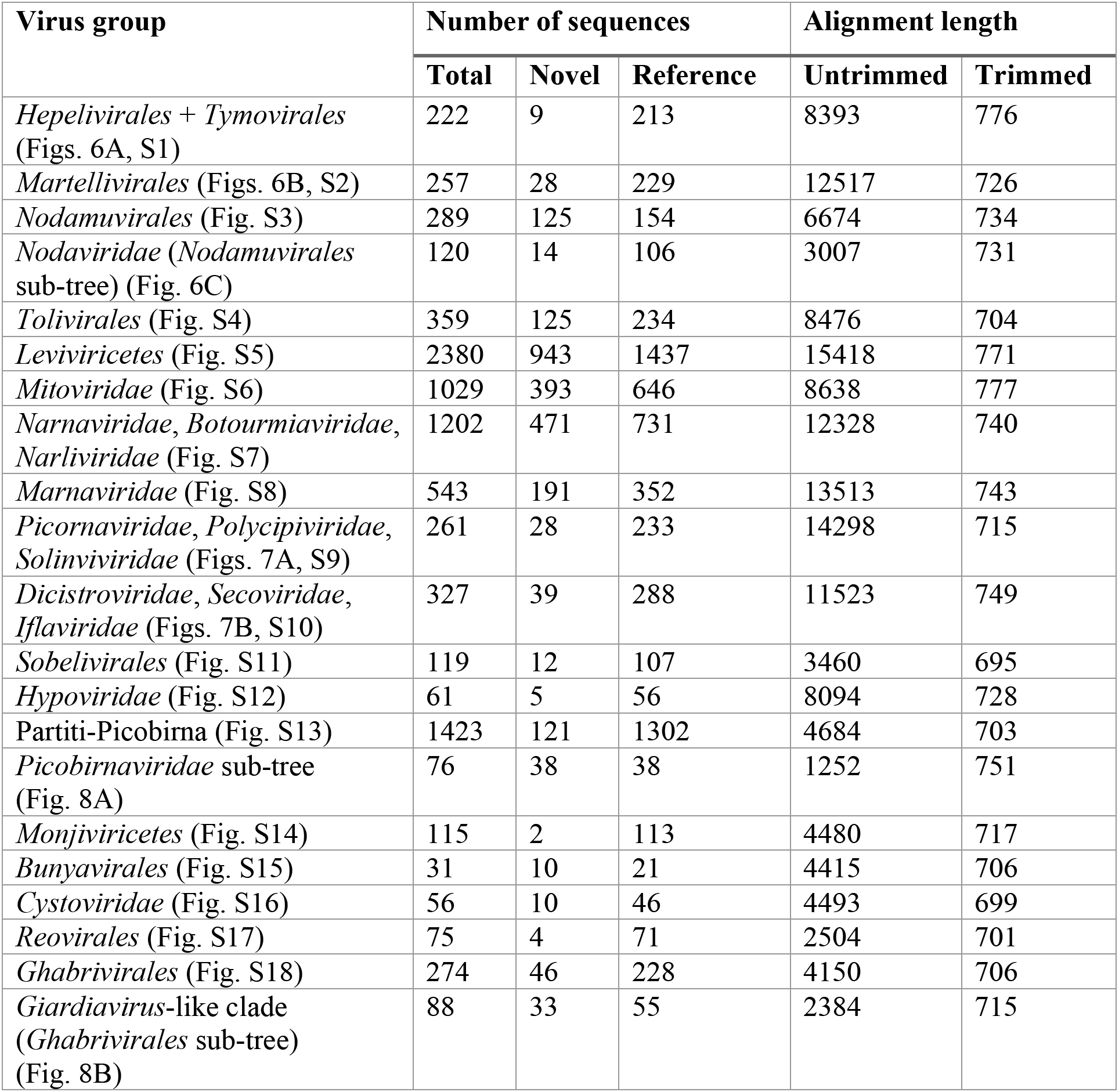
Sequence data sets used for phylogenetic analysis (Figures 6-8; Supplementary Figures 1-17).

| Virus group | Number of sequences |  |  | Alignment length |  |
| --- | --- | --- | --- | --- | --- |
|  | Total | Novel | Reference | Untrimmed | Trimmed |
| *Hepelivirales* + *Tymovirales* (Figs. 6A, S1) | 222 | 9 | 213 | 8393 | 776 |
| *Martellivirales* (Figs. 6B, S2) | 257 | 28 | 229 | 12517 | 726 |
| *Nodamuvirales* (Fig. S3) | 289 | 125 | 154 | 6674 | 734 |
| *Nodaviridae* (*Nodamuvirales* sub-tree) (Fig. 6C) | 120 | 14 | 106 | 3007 | 731 |
| *Tolivirales* (Fig. S4) | 359 | 125 | 234 | 8476 | 704 |
| *Leviviricetes* (Fig. S5) | 2380 | 943 | 1437 | 15418 | 771 |
| *Mitoviridae* (Fig. S6) | 1029 | 393 | 646 | 8638 | 777 |
| *Narnaviridae*, *Botourmiaviridae*, *Narliviridae* (Fig. S7) | 1202 | 471 | 731 | 12328 | 740 |
| *Marnaviridae* (Fig. S8) | 543 | 191 | 352 | 13513 | 743 |
| *Picornaviridae*, *Polycipiviridae*, *Solinviviridae* (Figs. 7A, S9) | 261 | 28 | 233 | 14298 | 715 |
| *Dicistroviridae*, *Secoviridae*, *Iflaviridae* (Figs. 7B, S10) | 327 | 39 | 288 | 11523 | 749 |
| *Sobelivirales* (Fig. S11) | 119 | 12 | 107 | 3460 | 695 |
| *Hypoviridae* (Fig. S12) | 61 | 5 | 56 | 8094 | 728 |
| Partiti-Picobirna (Fig. S13) | 1423 | 121 | 1302 | 4684 | 703 |
| *Picobirnaviridae* sub-tree (Fig. 8A) | 76 | 38 | 38 | 1252 | 751 |
| *Monjiviricetes* (Fig. S14) | 115 | 2 | 113 | 4480 | 717 |
| *Bunyavirales* (Fig. S15) | 31 | 10 | 21 | 4415 | 706 |
| *Cystoviridae* (Fig. S16) | 56 | 10 | 46 | 4493 | 699 |
| *Reovirales* (Fig. S17) | 75 | 4 | 71 | 2504 | 701 |
| *Ghabrivirales* (Fig. S18) | 274 | 46 | 228 | 4150 | 706 |
| *Giardiavirus*-like clade (*Ghabrivirales* sub-tree) (Fig. 8B) | 88 | 33 | 55 | 2384 | 715 |

Potentially novel virus sequences were aligned with members of the family, order, or multi-family clade of their respective closest DIAMOND BLASTX hit using MAFFT (v7.402) (Katoh and Standley, 2013). Sequence alignments were trimmed using trimAl (v1.4.1) (Capella-Gutierrez *et al*., 2009) to retain only the most conserved amino acid positions (between 695-777 residues in length) and to remove ambiguously aligned regions (see Table 1). Alignments were also visually assessed to identify and remove poorly aligned sequences. Maximum likelihood phylogenetic trees were then estimated on each of these alignments using IQ-TREE (v1.6.12) (Nguyen *et al*., 2015), employing the Le-Gascuel (LG) model of amino acid substitution – determined using ModelFinder within IQ-TREE – with 1,000 SH-aLRT replicates (Anisimova and Gascuel, 2006) to assess node support. All trees were visualised in R using packages ‘ape’ (v5.5) (Paradis and Schliep, 2019) and ‘ggtree’ (v3.0.2) (Yu *et al*., 2017). Probable host organisms for novel viruses were predicted based on the hosts of the established viruses with which they clustered most closely.

## 3. RESULTS

### 3.1 Data generation

We generated 26 sequencing libraries, 16 of which were from NSW farmland soil samples, with the remaining 10 from sediment taken from Denmark River in WA. Eighteen soil samples, representing three distinct soil types (vertosol, chromosol, and sodosol), two categories of land use (native vegetation or agricultural), and three depths (0-5 cm, 5-15 cm, and 15-30 cm), were taken from NSW farmland environments (Table S1). Vertosol, which is high in smectitic clay and has high agricultural potential, and chromosol, which has a loamy texture and moderate agricultural potential, were both collected from sites containing native vegetation or crops. Sodosol has a sandy surface texture with a high concentration of sodium and is nutrient poor. Due to the generally low capacity for crop growth, sodosol samples were taken from pasture and native vegetation sites. We were able to successfully extract RNA from six of these 18 samples, five of which we extracted in technical triplicates, totalling 16 RNA libraries for metatranscriptomic sequencing. No RNA was able to be extracted from vertosol or the 15-30 cm depths, and success in chromosol samples was restricted to the 0-5 cm depth despite multiple attempts on each sample.

The success of RNA extraction was similarly limited in the WA riverbank sediment samples. The 24 samples taken from a total of eight sites in the Swan-Canning Riverpark system (Perth, WA) were generally coarse and sandy in texture, while the 12 samples taken from four sites along Denmark River (Denmark, WA) were finer and muddier in comparison. No extractions from the Swan-Canning Riverpark system were successful, and RNA was only extracted from all three biological replicates in two of the four sites along Denmark River, for a total of 10 libraries able to be sequenced. Samples from which RNA was successfully extracted and sequenced are described in Table S1.

We generated approximately 2.68 billion paired-end reads from the 26 libraries successfully sequenced in this study, ranging from 66.7 million reads (native chromosol, NSW; library CN1A) to 198 million reads (native sodosol, NSW; library SN2A) per library. From these data, 6.7 million contigs were assembled, with contig numbers from individual libraries ranging from 90,913 (native sodosol, NSW; library SN2A) to 731,343 (riverbank sediment, WA; library KCAP1), with a median value of approximately 196,000.

### 3.2 RNA virome composition

Across the data set as a whole there were 12,292 contigs greater than 600 nucleotides (nt) in length that had DIAMOND BLASTX hits to sequences from the *Riboviria* (i.e., RNA viruses). However, a large proportion of contigs did not robustly align to reference sequences and could not be reliably assigned to any RNA virus taxa. Thus, a subset of 6,977 viral contigs was retained for analyses. The number of viral contigs per library ranged from 59 (native sodosol, NSW; SN2A) to 981 (riverbank sediment, WA; KCAP3) and the majority of contigs (5,209 out of 6,977) had less than 50% amino acid identity to reference sequences (Table S2). The sample with the lowest RNA viral abundance was NSW sodosol pasture (library SP2C) at 0.017%, while the highest abundance of 1.023% was found in WA riverbank sediment (library KCAP3) (Fig. 2A). Interestingly, the libraries with the lowest and highest diversity, MP2 and MP1, respectively, were both from the same WA riverbank site (Fig. 2B). While viral abundance differed significantly between sampling locations (p = 0.012), Shannon diversity did not (p = 0.458).

**Figure 2.**
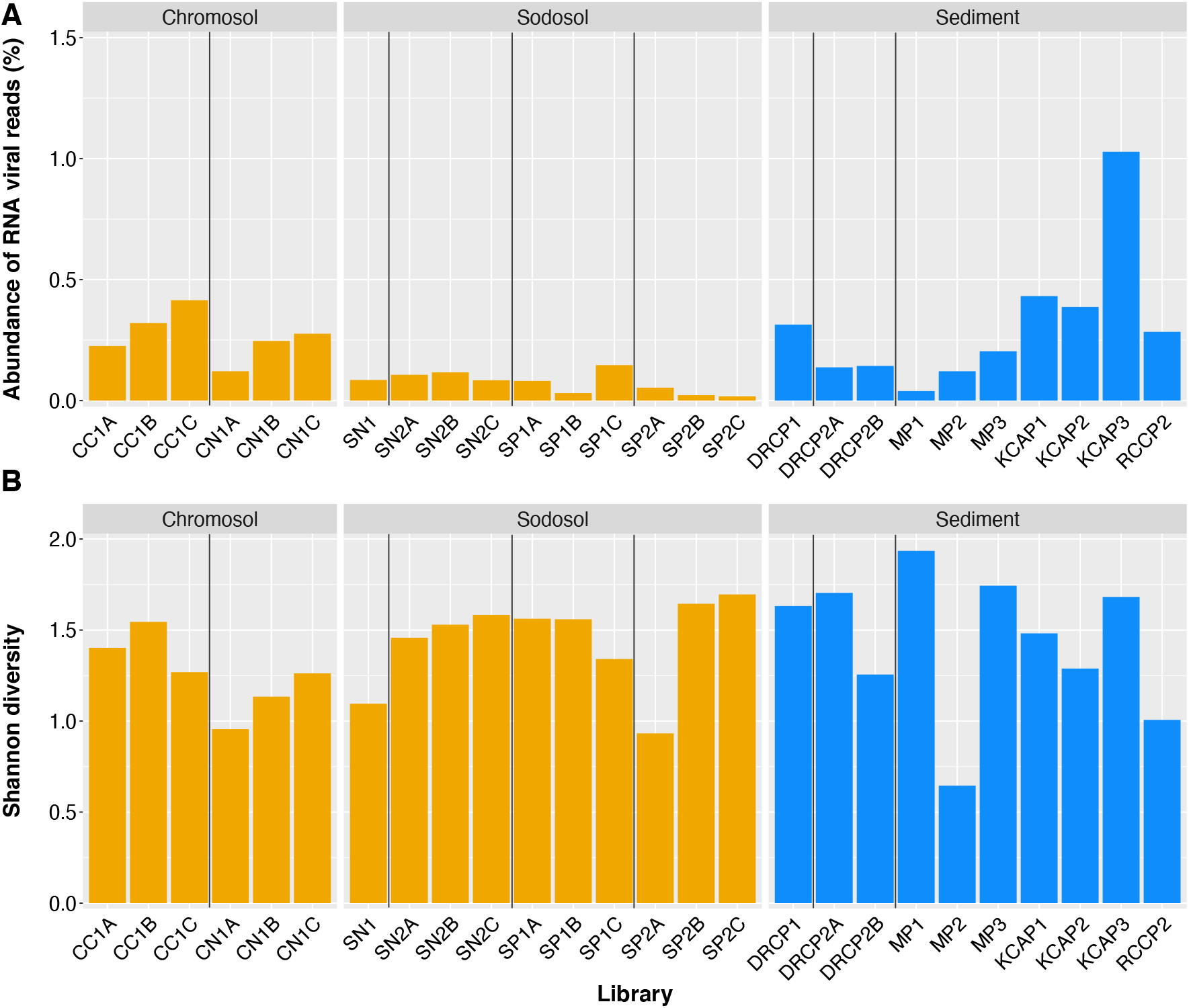
(A) Abundance of RNA virus reads as a proportion of the number of total reads and (B) RNA virus Shannon diversity indices of the meta-transcriptomic sequencing libraries generated from samples taken from NSW farmland soil (left and middle, in orange) and WA riverbank sediment (right, in blue). Letters (A, B, C) at the end of each library name indicate technical replicate extractions of the same sample and are grouped within vertical lines.

Sampling environment refers to the combination of soil type (chromosol, sodosol) and land use (native, cropping, pasture), which we found to be associated with soil virus abundance and richness (p < 0.001 in both cases) (Fig. 3A). To explore which factors were most likely to contribute to this effect, best-fit models were estimated considering soil type and land use separately. Only land use was included in the best fit model describing richness (p < 0.001), with agricultural soils (cropping and pasture) harbouring greater species richness than native soils in both chromosol and sodosol soil types (Fig. 3B). Soil type did not have a significant effect on richness (p = 0.58). Land use was also associated with viral abundance (p < 0.001), in which cropping soil had the highest abundance, pasture soil had the lowest, with native soils falling in the middle (Fig. 3B). However, it is difficult to determine if the agricultural purpose of the soil (i.e., cropping or pasture) was truly impacting viral abundance as soil type also had a significant influence on viral abundance (p < 0.001), which was higher in chromosol than in sodosol (Fig. 3C). As the cropping soils were only chromosol and the pasture soils were only sodosol, the pattern of abundance observed in Fig. 3C may have been due to the influence of soil type rather than agricultural purpose. However, while native chromosol had lower viral abundance than cropping chromosol, native sodosol had higher viral abundance than its counterpart used for pasture (Fig. 3A). This suggests an influence of both soil type and the purpose of agricultural land use, supported by the inclusion of both variables in the best-fit model for viral abundance. No ecological factors were associated with the Shannon and effective Shannon diversity indices (p = 0.14-0.61 and p = 0.10-0.51, respectively) (Fig. 3A-D). Finally, sampling depth did not significantly impact any abundance or alpha diversity indices of sodosol viruses (p = 0.13-0.79) (Fig. 3D), although it should be noted that soil depth affected our ability to extract quality RNA.

**Figure 3.**
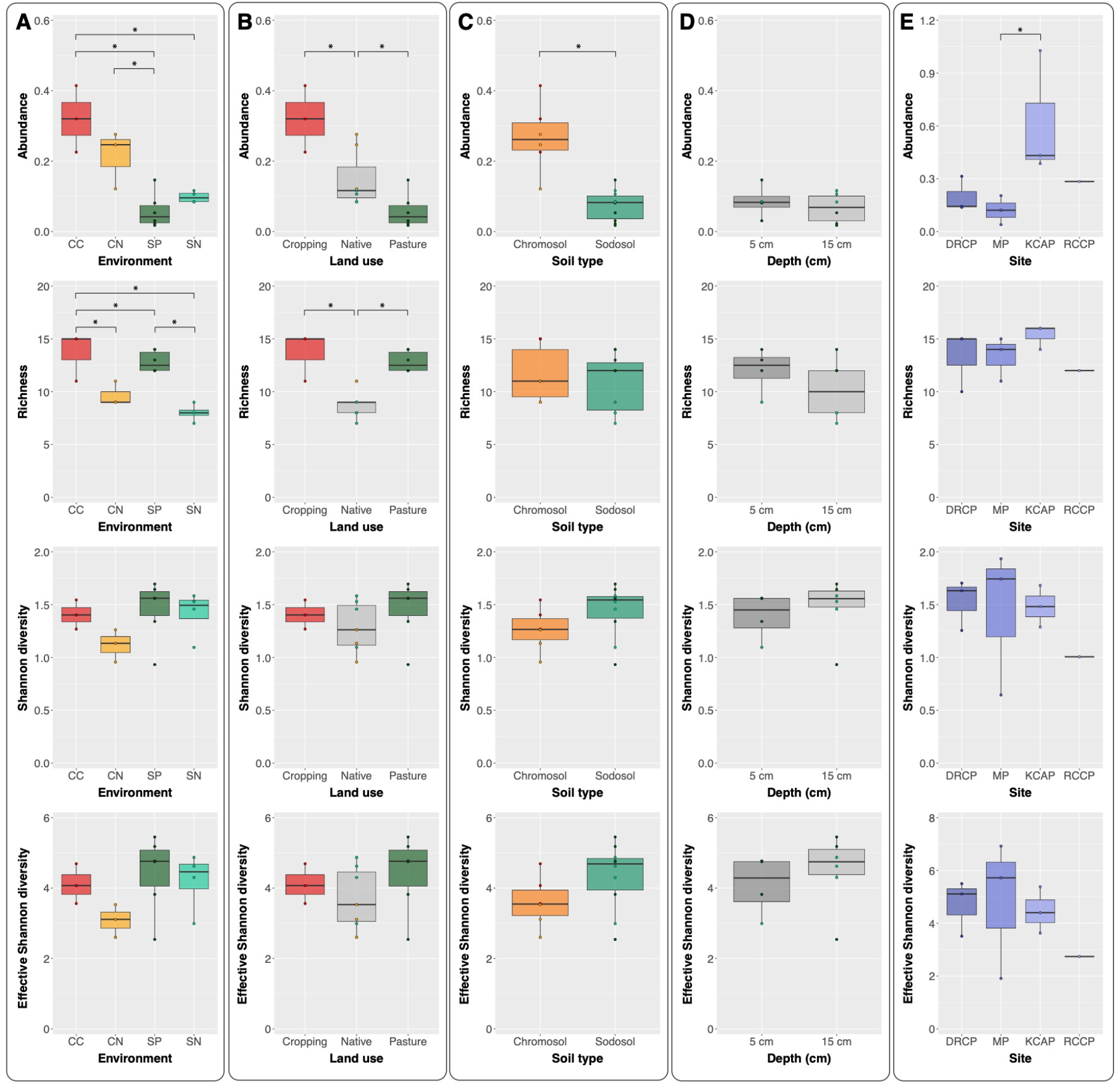
Abundance, richness, Shannon diversity, and effective Shannon diversity indices of NSW farmland soils plotted against (A) environment as a combination of soil type and land use, (B) soil type, (C) land use, (D) depth, and the same indices of WA riverbank sediments plotted against (E) sampling site. Asterisks indicate significant differences (p < 0.05) between pairs of ecological properties as determined by post hoc Tukey tests. Shorthand labels for environment indicate CC = cropping chromosol, CN = native chromosol, SP = pasture sodosol, and SN = native sodosol. Circles representing each library in columns B-D are coloured by sampling environment as per column A.

In the case of the sediment libraries, we measured the effect of sampling site on abundance, richness, Shannon diversity, and effective Shannon diversity (Fig. 3E). Only sampling site influenced viral abundance (p = 0.003), with libraries from the densely vegetated KCAP site having significantly higher abundance than the other sediment sites, and higher than any soil libraries (Fig. 2, Fig. 3E).

Relative virus abundances (i.e., the abundance of each viral group as a proportion of all *Riboviria* reads in each library) are shown in Fig. 4. The phylum *Lenarviricota* and orders *Tolivirales* and *Nodamuvirales* were present in all libraries, with the *Lenarviricota* generally comprising a large proportion, if not the majority, of reads. A notable exception was library DRCP2B (riverbank sediment, WA) where unclassified *Picornavirales* sequences comprised the majority of reads. As they were the most abundant groups, we classified the virome compositions of the phylum *Lenarviricota* and order *Picornavirales* to the class and family level, respectively (Fig. 4). In general, *Picornavirales* sequences comprised a greater proportion of reads in sediment libraries than in soil libraries. Furthermore, these sequences were mostly unclassified *Picornavirales* in sediment libraries, whereas in the soil libraries, the majority of *Picornavirales* sequences were from the family *Dicistroviridae*. Marnaviruses also appeared more frequently in the sediment libraries than in the soil libraries, likely due to their typically aquatic hosts, although pasture sodosol library SP1A had a high proportion of marnaviruses relative to the other *Picornavirales* groups and other soil libraries.

**Figure 4.**
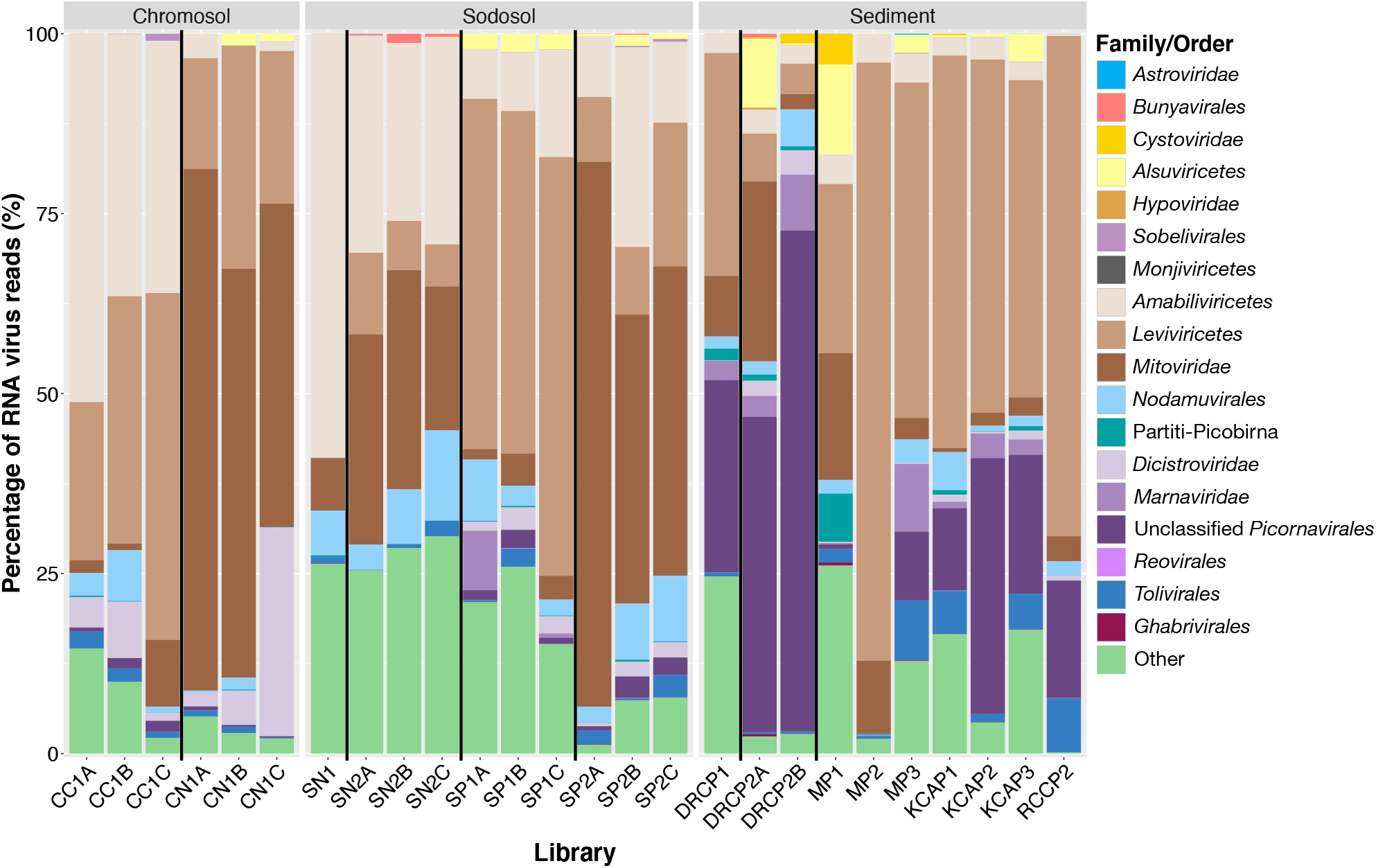
Virome compositions (family/order/phylum) for each sampling library. Relative proportions were determined by the total number of reads corresponding to contigs with DIAMOND BLASTX hits in each virus clade. Letters (A, B, C) at the end of each library name indicate technical replicate extractions of the same sample and are grouped within vertical lines. The more abundant phylum/order, *Lenarviricota* and *Picornavirales*, are further broken down into families or classes and are represented in shades of the same colour.

The *Lenarviricota* were abundant in all libraries. However, there was an overall decrease in the proportion of fungal-associated class *Amabiliviricetes* and family *Mitoviridae* in the sediment libraries compared to the soil libraries. Sediment libraries were instead dominated by bacteriophage of the class *Leviviricetes*, suggesting a switch in microbial community composition towards bacteria in sediment environments (Fig. 4). It is interesting to note that pasture sodosol samples taken 0-5 cm from the surface (libraries SP1A-C) were an exception to this trend, in which the *Leviviricetes* represented a much higher proportion of *Lenarviricota* sequences than seen in the other soil libraries (Fig. 4).

### 3.3 Phylogenetic analysis of novel virus species

We identified 2,562 novel virus RdRp sequences across all five RNA virus phyla, including those with positive-sense single-stranded RNA genomes (*Astroviridae, Picornavirales, Sobelivirales,* from the phylum *Pisuviricota*; *Alsuviricetes, Nodamuvirales,* and *Tolivirales* from the phylum *Kitrinoviricota*; and the phylum *Lenarviricota*), negative-sense single-stranded RNA genomes (*Bunyavirales* and *Monjiviricetes* from the phylum *Negarnaviricota*), and double-stranded RNA genomes (*Hypoviridae, Partitiviridae*, and *Picobirnaviridae* from the phylum *Pisuviricota*; and *Cystoviridae*, *Reovirales*, and *Ghabrivirales* from the phylum *Duplornaviricota*) (Table 1, Fig. 5, Supplementary Table 2). By predicting host organisms based on phylogenetic clustering, we suggest that viruses with bacterial, fungal, and protist hosts comprised 40.5%, 37.7%, and 8.3% of novel sequences, respectively, and those with plant and invertebrate hosts comprised 1.1% and 9.5% of novel sequences, respectively. We were unable to confidently assign a host organism for 2.9% of novel sequences, as these did not cluster closely enough to any reference sequences with a known host. Notably, no likely vertebrate-associated viruses were observed. The majority of novel sequences (1,807) fell into the microbe-associated phylum *Lenarviricota*. We now describe each of these groups in turn.

**Figure 5.**
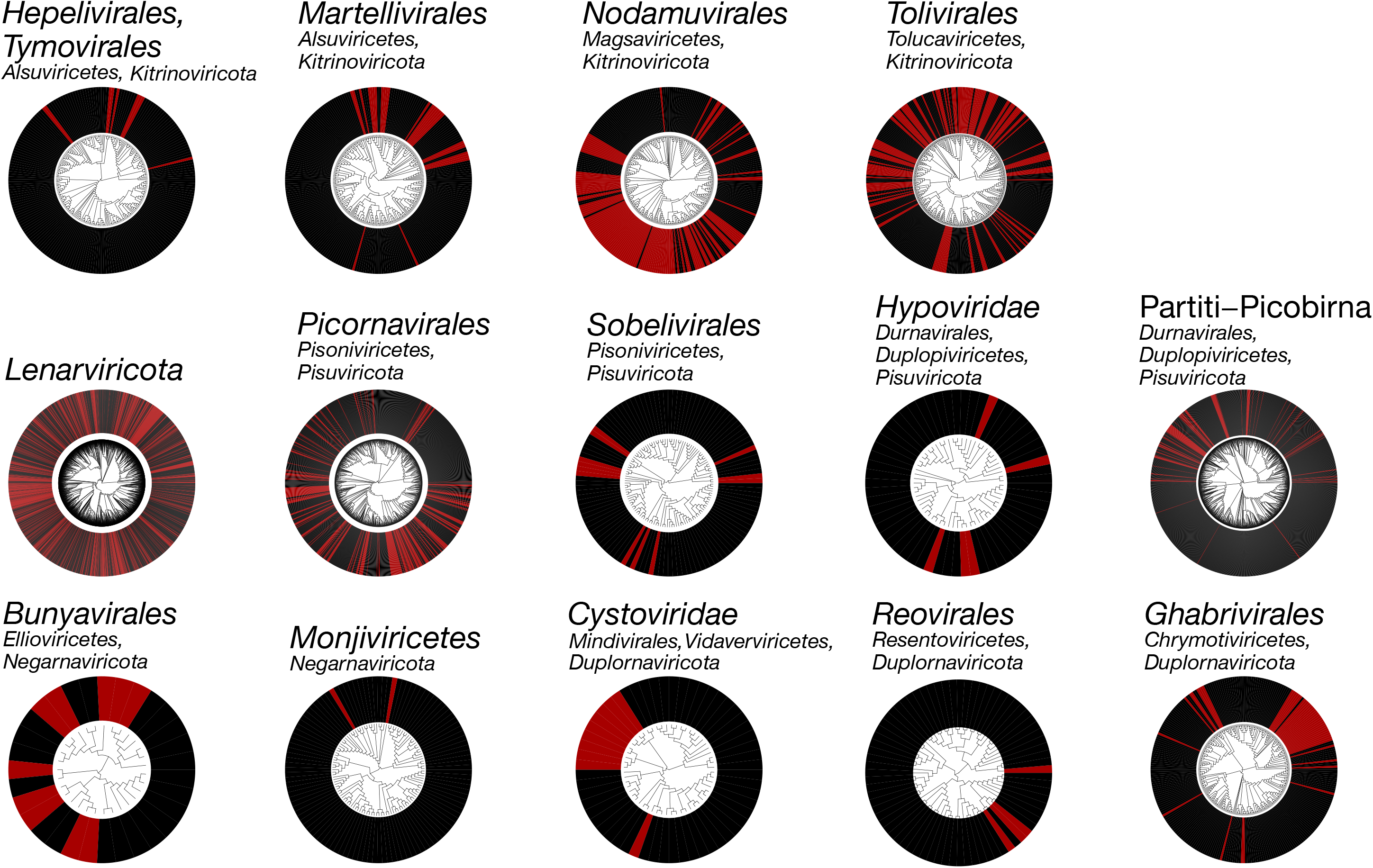
Phylogenetic trees of the RdRp protein sequences of viruses identified in this study, with reference sequences. Known viruses are shown in black, while putative novel viruses identified here are shown in red. All trees are midpoint rooted for clarity only. Individual phylogenetic trees for each clade are shown in Supplementary Figures 1–17. Details on the alignments used to estimate these phylogenies are shown in Table 1. Phylogenies of the phylum *Lenarviricota* and order *Picornavirales* were estimated for the purposes of visualising novel viruses but contain too much sequence divergence for robust alignments such that the phylogenies presented in the supplement are divided into multiple, smaller groups containing 1-3 related families each.

#### 3.3.1 Positive-stranded RNA viruses – phyla *Kitrinoviricota*, *Lenarviricota,* and Pisuviricota

##### Class Alsuviricetes (Kitrinoviricota) and bastroviruses (Pisuviricota)

Five novel sequences clustered within the order *Tymovirales* (Fig. 6A, Supplementary Fig. 1). Two appeared to be novel marafiviruses (*Tymoviridae*), while the other three clustered with the divergent mycoflexivirus Botrytis virus F (*Gammaflexiviridae*). A single virus clustered within the *Hepelivirales*. All six novel viruses from the *Tymovirales* and *Hepelivirales* were identified from riverbank sediment samples.

**Figure 6.**
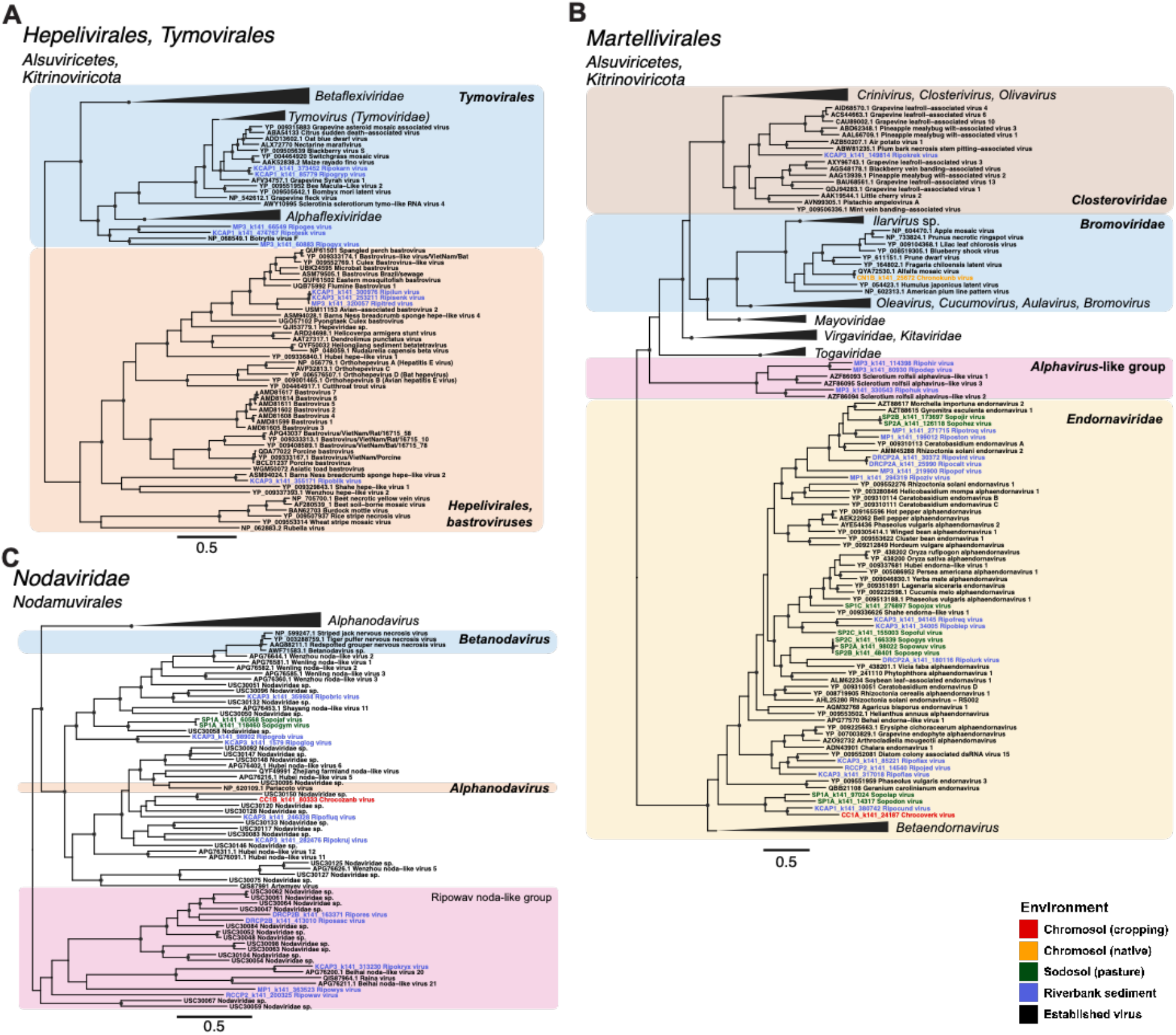
Phylogenetic trees of RdRp sequences from (A) the orders *Hepelivirales* and *Tymovirales*, (B) the order *Martellivirales*, and (C) the family *Nodaviridae*. Known viruses are shown in black, while putative novel viruses identified here are coloured by sampling environment. Trees are midpoint rooted for clarity only and branch lengths are scaled according to the number of amino acid substitutions per site. Black circles represent node support ≥80% using 1,000 SH-aLRT replicates. Phylogenies with collapsed branches expanded are shown in Supplementary Figures 1-3.

Despite being classified as members of the *Astroviridae*, bastroviruses encode a *Hepeviridae*-like RdRp protein due to a recombination event (Oude Munnink *et al*., 2016). Three novel bastrovirus-like sequences were identified in riverbank sediment, forming a divergent sister group to avian-associated bastrovirus 2 (USM11153) (Fig. 6A, Supplementary Fig. 1). Given the broad host range of bastroviruses, the host of these novel bastroviruses cannot be confidently inferred.

Several novel sequences were identified in the order *Martellivirales* (Fig. 6B, Supplementary Fig. 2). These included an ampelovirus (*Closteroviridae*), an ilarvirus (*Bromoviridae*), and 19 alphaendornaviruses (*Endornaviridae*). Notably, three novel viruses – Ripohir virus, Ripodep virus, and Ripohuk virus – had 27-31% sequence similarity to and clustered with three divergent mycoviruses – *Sclerotium rolfsii* alphavirus-like virus 1-3 (AZF86093-5). This clade fell as a sister-group to the plant-infecting families *Closteroviridae*, *Bromoviridae*, *Mayoviridae*, *Virgaviridae*, *Kitaviridae*, and the animal-infecting, insect-borne *Togaviridae*.

##### Nodamuvirales (Kitrinoviricota)

Our phylogenetic analysis of the *Nodaviridae* revealed a conflict with the current taxonomy (Fig. 6C). A clade including the alphanodavirus species Pariacoto virus (NP_620109) formed a sister group to the genus *Betanodavirus* rather than clustering with the remaining alphanodaviruses. This clade comprised several environmental noda-like viruses, including the novel noda-like viruses Ripofluq virus, Chrocozanb virus, and Ripokruj virus, and may represent the existence of a third genus within the *Nodaviridae*.

A total of nine viruses fell within the *Nodaviridae*, and another five sequences fell within a group of noda-like viruses forming a sister clade to the *Nodaviridae* (Fig 6C, Supplementary Fig. 3). This sister clade – provisionally named Ripowav noda-like group – may represent a fourth genus within the *Nodaviridae*. However, the largest expansion of virus diversity was in a sister clade to the *Nodaviridae* that likely comprises a new family. This putative family included Lake Sinai virus 1 and 2 (*Sinhaliviridae*), many noda-like viruses previously identified from soil (Chen *et al.,* 2022) and invertebrates (Shi *et al*., 2016) in China, and novel viruses from all environments sampled (Supplementary Fig. 3).

##### Tolivirales (Kitrinoviricota)

Novel *Tolivirales* sequences were also identified in all environments sampled, although those clustering within the plant-associated *Tombusviridae* were predominantly identified in riverbank sediment libraries (Supplementary Fig. 4). A total of 29 sequences (27 from riverbank sediments) formed sister lineages to the subfamilies *Procedovirinae* and *Calvusvirinae*, and 12 formed sister lineages to the entire family *Tombusviridae*.

All sampling environments were represented in the 20 novel sequences clustering within a clade of diverse tombus-like viruses (Supplementary Fig. 4). This clade also included the sole species within the *Carmotetraviridae* - Providence virus (AMQ67162) - a unique virus isolated from arthropod (lepidopteran) tissue that is also capable of replicating in plant and mammalian cell lines (Jiwaji *et al*., 2019). Finally, 38 novel sequences fell into the most divergent clade in this phylogeny, which largely comprised previously identified environmental viruses, as well as some sourced from invertebrates and animal faecal samples (Supplementary Fig. 4).

##### Lenarviricota

A remarkable 1,807 novel sequences were identified within the phylum *Lenarviricota*, comprising the majority of novel viruses found in this study. Novel sequences were sourced from sediment and both farmland soil types and land uses. Due to the high level of sequence divergence across the *Lenarviricota*, phylogenetic trees were estimated on sub-alignments of (i) the class *Leviviricetes* (i.e., bacteriophage; Supplementary Fig. 5), (ii) the family *Mitoviridae* (Supplementary Fig. 6), and (iii) the families *Narnaviridae*, *Botourmiaviridae* and the newly proposed *Narliviridae* (Supplementary Fig. 7).

In total, 943 novel leviviruses were identified and every sampling environment was represented in this group of viruses. In many large clades, the majority of viruses were novel sequences identified in this study (Supplementary Fig. 5). Similarly, several clades mainly comprising novel mitoviruses were identified in soil environments. Of particular interest was the clade of seven native chromosol, five pasture sodosol, and two sediment viruses that clustered with a mitovirus - Kinsystermes vitus (QQM15243) - identified in a termite, although their use of the mitochondrial genetic code suggests they likely infect the fungal hosts that are typical of mitoviruses (Supplementary Fig. 6). Eight sediment and two pasture sodosol mitoviruses formed a sister clade to viruses likely associated with the microbiomes of arthropods and vertebrates, although as these viruses were obtained from metagenomic studies their true host association is unclear.

In the *Narnaviridae-Botourmiaviridae-Narliviridae* phylogeny multiple new, distinct clades comprising viruses from different sampling sources were observed, typically biased towards sediment or soil environments. These novel viruses typically shared <50% amino acid identity with any reference sequence. One clade of 124 entirely new viruses was identified within the *Narliviridae*, potentially representing a genus within this family (Supplementary Fig. 7). The viruses comprising this group included 30 from sediment, 27 and 17 from pasture and native sodosol, respectively, and 42 and 8 from cropping and native chromosol, respectively (Supplementary Fig. 7). Most of the novel *Botourmiaviridae* identified here were associated with soil samples. This contrasted with most of the viruses detected from other families within the *Lenarviricota* that were predominantly associated with sediments (Supplementary Fig. 7). Indeed, only three of the 62 novel botourmiaviruses were identified in sediment, and only from a single sediment library (MP3). A tendency for soil over sediment has previously been observed in the *Botourmiaviridae* (Chen *et al*., 2022). We also identified 35 novel *Narnaviridae* species in sediment, 10 and 8 in pasture and native sodosol, respectively, and 19 and 7 in cropping and native chromosol, respectively. Notably, the majority of the novel viruses identified in the *Lenarviricota* occurred in lineages that were divergent from known viral families, including viruses from all five sampling environments (Supplementary Figs. 5-7).

##### Picornavirales (Pisuviricota)

The main expansion of novel sequences in this order was in the *Marnaviridae* and *Dicistroviridae,* as well as several unclassified *Picornavirales* species that fell outside of defined families. Again, the scale of novel diversity in this order warranted splitting it into smaller groups of 1-3 families for robust phylogenetic analysis: (i) the *Marnaviridae* (Supplementary Fig. 8), (ii) the *Picornaviridae*, *Polycipiviridae*, *Solinviviridae* (Fig. 7A, Supplementary Fig. 9), and (iii) the *Dicistroviridae*, *Secoviridae*, *Iflaviridae* (Fig. 7B, Supplementary Fig. 10). Each phylogeny also included several clades of unclassified *Picornavirales*.

**Figure 7.**
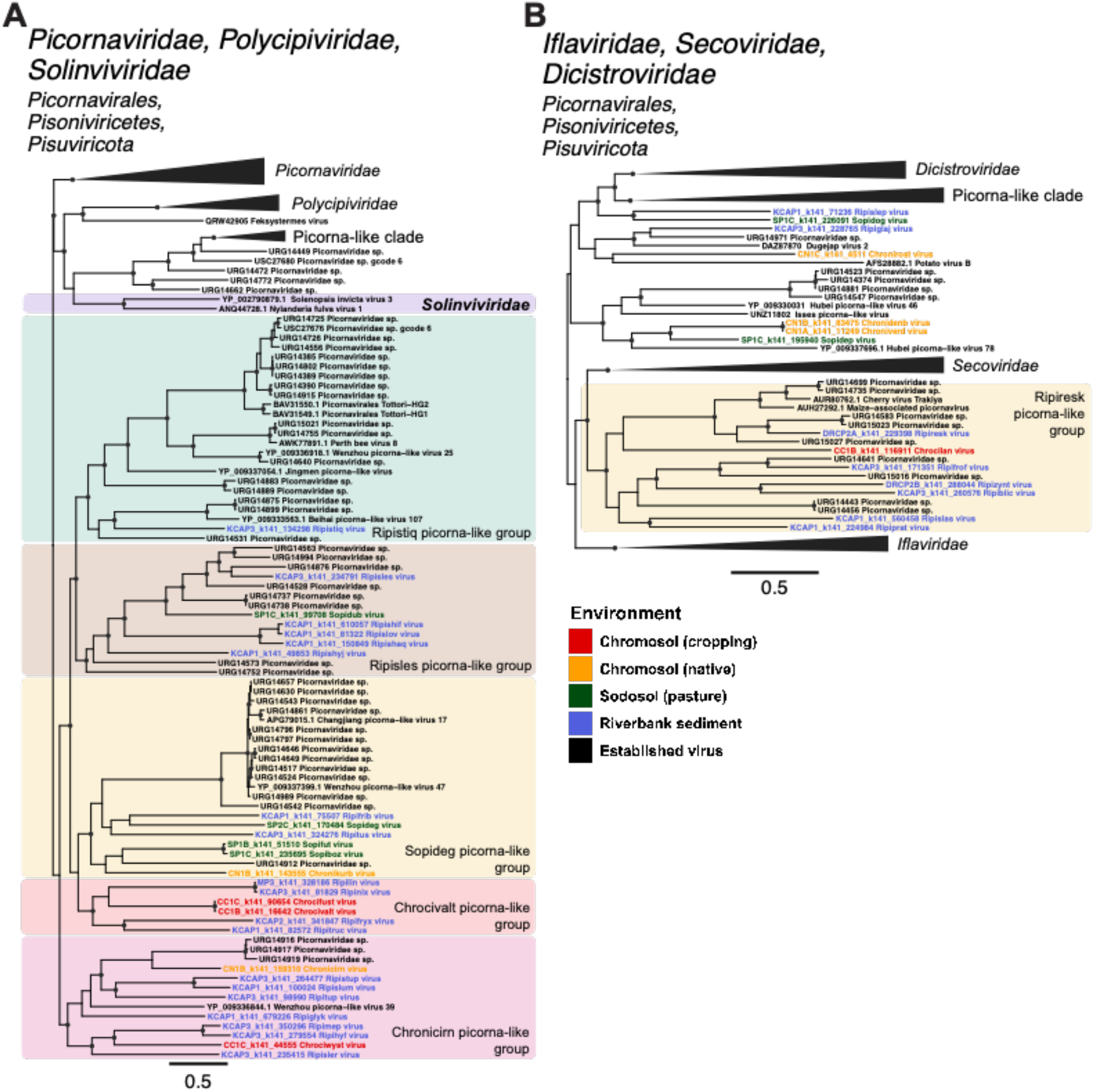
Phylogenetic trees of the RdRp sequences from the order *Picornavirales*, displaying the expansion of novel, unclassified picorna-like clades surrounding (A) the *Picornaviridae*, *Polycipiviridae*, *Solinviviridae* and a clade of previously identified picorna-like sequences, and (B) the *Dicistroviridae*, *Iflaviridae*, and *Secoviridae*. Known viruses are shown in black, while putative novel viruses identified here are coloured by the sampling environment. Trees are midpoint rooted with branch lengths scaled according to the number of amino acid substitutions per site. Black circles represent node support ≥80% using 1,000 SH-aLRT replicates. Phylogenetic trees with collapsed branches expanded are shown in Supplementary Figures 9 and 10.

A total of 191 novel viruses with sequence similarity (22-98%) to the *Marnaviridae* were identified. The majority were sourced from riverbank sediment, which is unsurprising given that marnaviruses are typically associated with marine organisms and environments. At the genus level, 17 viruses from sediment clustered with the sogarnaviruses, 33 with the salisharnaviruses, and one with the *bacillarnaviruses* (Supplementary Fig. 8). Eleven sequences clustering in the genus *Marnavirus* were also identified in sediment, grouping with Antarctic picorna-like virus 3 (AKG93964) and *Picornaviridae* sp. (URG14782, URG14392, URG14815). These sequences formed a sister clade to the group containing the sole member of the genus *Marnavirus* - Heterosigma akashiwo RNA virus (YP_009047193) (Supplementary Fig. 8). Novel sequences from soil and sediment environments were also identified in the genera *Kusarnavirus*, *Locarnavirus*, and *Labyrnavirus*. Finally, several unclassified virus lineages fell between certain genera: four clades between *Bacillarnavirus* and *Marnavirus*, three between *Kusarnavirus* and *Locarnavirus*, and three falling basal to all genera except for *Labyrnavirus* (Supplementary Fig. 8).

A group of 28 novel picorna-like viruses fell in a divergent sister clade to the established families *Picornaviridae*, *Polycipiviridae*, and *Solinviviridae* (Fig. 7A., Supplementary Fig. 9). The majority of published sequences in this sister clade were metagenomically sourced from environmental or invertebrate samples. The diversity within this clade suggests that the creation of several new virus families may be warranted, with five well-supported clusters able to be identified (Fig. 7A, Supplementary Fig. 9). Eighteen novel viruses were identified in the *Dicistroviridae*, and a further fourteen novel picorna-like viruses fell in lineages of diverse, unclassified *Picornavirales* that fell as sister lineages to the *Dicistroviridae* (Supplementary Fig. 10). Another clade of interest – provisionally named the Ripiresk picorna-like group – fell between the plant-infecting *Secoviridae* and insect-associated *Iflaviridae* (Fig. 7B, Supplementary Fig. 10). This group included six novel viruses from sediment, one from cropping chromosol, several previously identified environmental picorna-like viruses, and two picornaviruses identified in plants (Fig. 7B, Supplementary Fig. 10).

##### Sobelivirales (Pisuviricota)

Three viruses fell within the fungi-associated family *Barnaviridae*, while another two sequences formed a clade with the sole member of the dinoflagellate-associated *Alvernaviridae* and three divergent sobemo-like viruses (Supplementary Fig. 11). Several other sobeli-like viruses fell in clades of unclassified *Sobeliviridae*. Of note, were three novel soil – Sonifin virus, Sopibym virus, and Sopibym virus – which formed a cluster with solemoviruses predominantly associated with insects including termites, rabbit fleas, and flies (Supplementary Fig. 11).

##### Hypoviridae (Pisuviricota)

Four novel *Hypoviridae* sequences were identified in sediment and one in cropping chromosol. These sequences were highly divergent and had low node support, such that their true phylogenetic placement could not be robustly determined (Supplementary Fig. 12).

##### Partiti-Picobirnaviruses (*Pisuviricota*)

Novel sequences were identified in both the *Partitiviridae* and *Picobirnaviridae* (Supplementary Fig. 13). The majority of novel picobirnaviruses were found in divergent, unclassified clades. The largest cluster of novel sequences were sampled from riverbank sediment and grouped with a clade including termite microbiome-associated picobirnaviruses identified by Le Lay *et al*. (2020) (Fig. 8A). Of these, 20 novel sediment picobirnaviruses formed a single monophyletic group.

**Figure 8.**
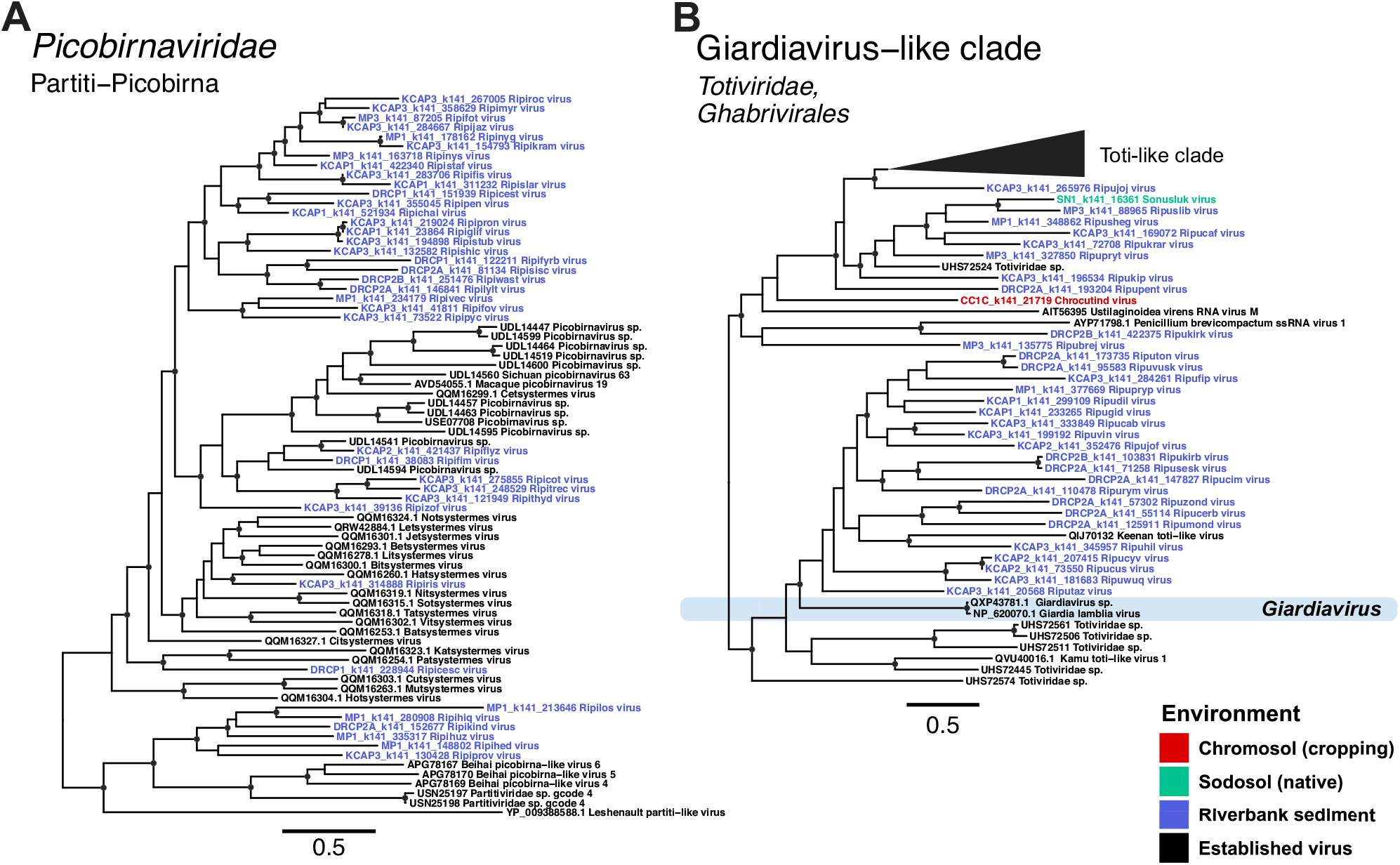
Phylogenetic trees of RdRp sequences from (A) a subset of the *Picobirnaviridae*, and (B) a clade of *Giardiavirus*-like sequences from the *Totiviridae*. Known viruses are shown in black, while putative novel viruses identified here are coloured by the sampling environment. Trees are midpoint rooted with branch lengths scaled according to the number of amino acid substitutions per site. Black circles represent node support ≥80% using 1,000 SH-aLRT replicates. Phylogenetic trees with collapsed branches expanded are shown in Supplementary Figures 13 and 17.

#### 3.3.2 Negative-stranded RNA virus families (*Negarnaviricota*)

It is striking that only twelve putative novel species of negative-sense RNA viruses were identified, two of which fell within the class *Monjiviricetes*. These viruses – Ripivex virus and Ripivax virus - were identified from a single riverbank sediment library (MP3). Ripivex virus clustered within the family *Mymonaviridae* (although with low bootstrap support), while Ripivax fell basal to the family (Supplementary Fig. 14).

The remaining ten negative-stranded RNA viruses fell in two clades of plant pathogenic fungi-associated *Bunyavirales*. The first clade, denoted the Sonekey bunya-like group, included four sequences related to bunyaviruses found in grapevine downy mildew lesions caused by *Plasmopara viticola* (Supplementary Fig. 15). In the second clade, the Soperolo bunya-like group, five sequences clustered with bunyaviruses associated with *Phytophthora cactorum* and *Halophyophthora* species. The final sequence, Chrocemuse virus, fell between the Sonekey and Soperolo bunya-like groups (Supplementary Fig. 15). It is interesting that five of the six viruses in Soperolo bunya-like group were identified in soil, given that several of their relatives are associated with predominantly marine-inhabiting host organisms (*Halophytophthora* species).

#### 3.3.3 Double-stranded RNA virus families (*Duplornaviricota*)

##### Cystoviridae

All ten novel cysto-like viruses were identified from WA riverbank sediment samples. One sequence (Ripucork virus) fell within a clade including the formally ratified species *Cystovirus phi13*, *Cystovirus phiYY*, and *Cystovirus phi6*. The remaining nine sequences formed two intermediate lineages between the two major clades in this family and clustered by sampling site (Supplementary Fig. 16). The first clade comprised five viruses identified in libraries from sediment samples taken further inland than the other four viruses, which, along with Jiangsu sediment cystovirus (QYF49681) and *Cystovirus phi8* (NP_524561), comprised the second clade.

##### Reovirales

One divergent reo-like virus in sodosol pasture (denoted Soputhoc virus) grouped with Hubei reo-like virus 10 and 11 (APG79149 and APG79051), falling as a sister lineage to the genus *Rotavirus* (a vertebrate-associated genus) in the *Sedoreoviridae*, although with low node support (Supplementary Fig. 17). The three other novel reo-like sequences (one identified in sodosol pasture, the others in riverbank sediment) fell in a small sister clade to the genus *Fijivirus* within the family *Spinareoviridae* (Supplementary Fig. 17).

##### Ghabrivirales

Novel species in this order were identified predominantly in sediment, with a small number from both sodosol environments and cropping chromosol (Supplementary Fig. 18). Of note, 22 novel toti-like sequences sourced from sediment greatly expanded a clade that previously only included Giardia lamblia virus (NP_620070), Giardiavirus sp. (QXP43781), and Keenan toti-like virus (QIJ70132) (Fig. 8B). This may represent an expansion of the *Totiviridae* genus *Giardiavirus*, which currently contains only one accepted species – Giardia lamblia virus. Other novel species in the *Totiviridae* included a victori-like and a totivirus from cropping chromosol and pasture sodosol, respectively, as well as two sequences identified from sediment samples that clustered with one of several lineages of unclassified *Totiviridae* (Supplementary Fig. 18). A novel alphachryso-like virus (Riputesc virus) and a divergent megabirna-like virus (Ripusarb virus) were also identified in sediment (Supplementary Fig. 18).

## DISCUSSION

Numerous novel RNA viruses have been identified in Australia’s unique ecosystems (Geoghegan *et al*., 2021; Harvey *et al*., 2018; Le Lay *et al*., 2020; Mahar *et al*., 2020; Mu *et al*., 2018; Pyke *et al*., 2021; Chang *et al*., 2021; Van Brussel *et al*., 2022; Wille *et al*., 2018). While efforts have been made to characterise the overall microbial communities of Australian soils (Bowd *et al*., 2022; Xue *et al*., 2022; Pino *et al*., 2023), little work has been done on the Australian soil virome. We generated meta-transcriptomic data on 26 libraries from 16 soil and sediment samples taken from eastern (NSW) and western (WA) Australia. From these, we identified a remarkable 2,562 novel viral RdRp sequences across all five RNA virus phyla, of which 1,807 belonged to the phylum *Lenarviricota*, classically associated with microbial species. The discovery of 2,562 putative viruses from such a small number of meta-transcriptomic sequence libraries showcases the extensive, untapped diversity of RNA viruses in Australian soil and sediment environments. Viruses were detected across 15 viral orders and, in many cases, were so diverse that they would constitute new viral genera and even families.

A relationship between virome composition and land use has been previously observed for both DNA viruses (Narr *et al*., 2017; Liao *et al*., 2022) and RNA viruses (Hillary *et al*., 2022) in soil on other continents. Likewise, local sampling environment has been shown to be associated with viral abundance and diversity (Chen *et al*., 2022; Durham *et al*., 2022). We found land use and sampling environment to be significantly associated with the abundance and richness of RNA viruses in Australian soils. However, more specific soil factors such as pH and soil nutrient levels have been identified as determinants of viral abundance and diversity (Narr *et al*., 2017; Chen *et al*., 2022; Liao *et al*., 2022). These factors also determine the community compositions of soil-dwelling microbial hosts (Wang *et al*., 2019; He *et al*., 2022), and are therefore likely play a role in the viral abundance and diversity trends observed in this study. Revealing the precise relationships between Australian soil ecosystems and the viruses within them will not only rely on expansive sampling of diverse environments, but also the generation of thorough ecological metadata.

Despite a small sample size, we detected viruses that spanned the entire diversity of the *Lenarviricota*. Such remarkable genetic diversity, as well as the presence of bacterial- and eukaryote-associated families (Hillman and Cai, 2013) and an often very simple genome structure, suggests that this phylum may comprise the oldest extant RNA viruses. Meta-transcriptomic studies of diverse ecosystems have consistently detected members of the *Lenarviricota*, such that the diversity of this phylum has greatly increased in recent years. Despite their microbial association, these viruses have been detected in studies of both vertebrates (Mahar *et al*., 2020; Wille *et al*., 2020) and invertebrates (Shi *et al*., 2016; Kondo *et al*., 2020; Le Lay *et al*., 2020; Thongsripong *et al*., 2021). Although they are unlikely to be infecting these animals directly, it is clear that these highly diverse viruses are present in virtually all environments (Starr *et al*., 2019; Wolf *et al*., 2020; Chen *et al*., 2022; Neri *et al*., 2022). Hence, it is no surprise that 1,807 novel *Lenarviricota* sequences were identified from both soil types and land uses sampled here, as well as a considerable number from riverbank sediment.

Within the *Lenarviricota*, the *Narnaviridae* and *Botourmiaviridae* are currently placed in different virus classes but have been proposed to comprise a single taxonomic class along with the newly proposed *Narliviridae* (Sadiq *et al*., 2022). The phylogeny generated here, with an additional 1,807 viral sequences expanding the known *Lenarviricota* diversity, supports the proposal of Sadiq *et al*. (2022) that the *Narliviridae* and *Botourmiaviridae* form sister clades to the exclusion of the *Narnaviridae*. Furthermore, the genus *Ourmiavirus* did not fall within the *Botourmiaviridae*, where it is currently classified, but instead clustered with the *Narliviridae*. This corroborates previous findings that these three ourmiaviruses are more closely related to the narliviruses than the botourmiaviruses (Sadiq *et al*., 2022).

Based on the nature of the samples (soil and sediment) and the known host ranges of families with which these novel viruses share sequence similarity, the majority of host taxa evaluated in this study were most likely associated with plant or microbial rather animal hosts. The limited proportion of viruses that were predicted to have animal hosts (less than 10%) were also most likely associated with invertebrates such as insects rather than vertebrates. The novel mitoviruses identified here appeared to utilise the mitochondrial genetic code, which changes the UGA codon from a stop codon to Tryptophan and is typical of fungi-infecting mitovirus genomes, suggesting that these mitoviruses also infect fungi (Cole *et al*., 2000). As the *Leviviricetes* and Cystoviridae are a class and family of bacteriophage (King *et* al., 2013; Poranen *et al*., 2017), the 953 novel viruses within them are also likely bacteriophage. The detection of large numbers of picobirnaviruses in environmental samples supports the hypothesis that these viruses do not infect vertebrate hosts (as is routinely assumed when associated with vertebrate faecal samples [Malik *et al*., 2014; Delmas *et al*., 2019]), but are instead associated with microflora present in animal gastrointestinal systems or components of their diet (Ghosh and Malik, 2021). Novel species in the *Tombusviridae* were likely to be associated with plants due to their phylogenetic proximity to established plant viruses. Similarly, the four novel viruses that clustered within the established dicistroviruses are likely to be insect-associated. Therefore, while the metagenomic nature of this study makes it inherently difficult to confidently determine the hosts of the novel viruses identified, assuming similar host ranges to their closest phylogenetic relatives reveals a remarkable variety of potential host organisms ranging from microbes to plants to invertebrates. Hence, viruses are infecting the majority, if not all, key players in the ecological processes of terrestrial and aquatic soil systems.

A limitation of this study was the initial extraction of genetic material. Of 90 potential sequencing libraries (including technical triplicates for the NSW farmland soil samples), RNA was only successfully extracted from 26 samples at low but detectable concentrations of 0.26-10 ng/µL. This prevented us from conducting robust statistical analyses on the various ecological factors that might shape RNA viral abundance and diversity. The inability to extract RNA from any samples collected 15-30 cm from the soil surface may be a result of the reduction in host organisms at this depth as deeper layers of soil have been shown to harbour limited microbial diversity compared to surface layers (Hao *et al*., 2021; Zhao *et al*., 2021). We were also unable to extract RNA from the vertosol, even from surface level soil which was successful in both chromosol and sodosol. This is likely due to the high clay content of vertosol, which has a negative effect on RNA yield under a variety of extraction protocols (Novinscak and Fillon, 2011). Yields may have also been impacted due to limitations in sample preservation, indicating a clear need for a more refined sampling and storage cold chain to effectively extract RNA from remote soil and sediment environments.

The discovery of 2,562 novel viruses spanning all five RNA viral phyla and a potential host range of bacteria, protists, fungi, plants, and invertebrates shows that Australian terrestrial environments are evidently an untapped resource for RNA virus diversity. These environments may harbour entire families of ecological and evolutionary importance, likely reflecting the vast array of flora and fauna that is unique to the continent. Our work provides an initial view of the Australian terrestrial RNA virosphere, as well as the broad environmental properties such as land use and soil type that may be driving viral composition.

## Data Availability

The raw sequence read data for this study are available in the NCBI Sequence Read Archive (SRA) database under the BioProject PRJNA981585 and SRA accession numbers SRR26298330-SRR2629835. All genome sequences used in the phylogenetic analysis are available in NCBI GenBank under the accession numbers XXXX-YYYY.

## CRediT authorship contribution statement

Sabrina Sadiq: Conceptualization, Formal analysis, Investigation, Methodology, Writing – original draft, Writing – review & editing, Visualization.

Erin Harvey: Investigation, Writing – review & editing.

Jonathon C. O. Mifsud: Investigation, Writing – review & editing.

Budiman Minasny: Data curation, Writing – review & editing.

Alex. B. McBratney: Data curation, Writing – review & editing.

Liana E. Pozza: Data curation, Investigation, Writing – review & editing.

Jackie E. Mahar: Investigation, Writing – original draft, Writing – review & editing.

Edward C. Holmes: Conceptualization, Resources, Writing – original draft, Writing – review & editing.

## Acknowledgments

We thank Dr Michelle Wille for previously providing the modified Rhea alpha diversity script sets. We also thank Dr Edward Jones for assisting with sample collection at the Nowley Farm sites.

