## Supplemental Figures for "Australian terrestrial environments harbour extensive RNA virus diversity"

### *Hepelivirales, Tymovirales*

*Alsuviricetes,*
*Kitrinoviricota*

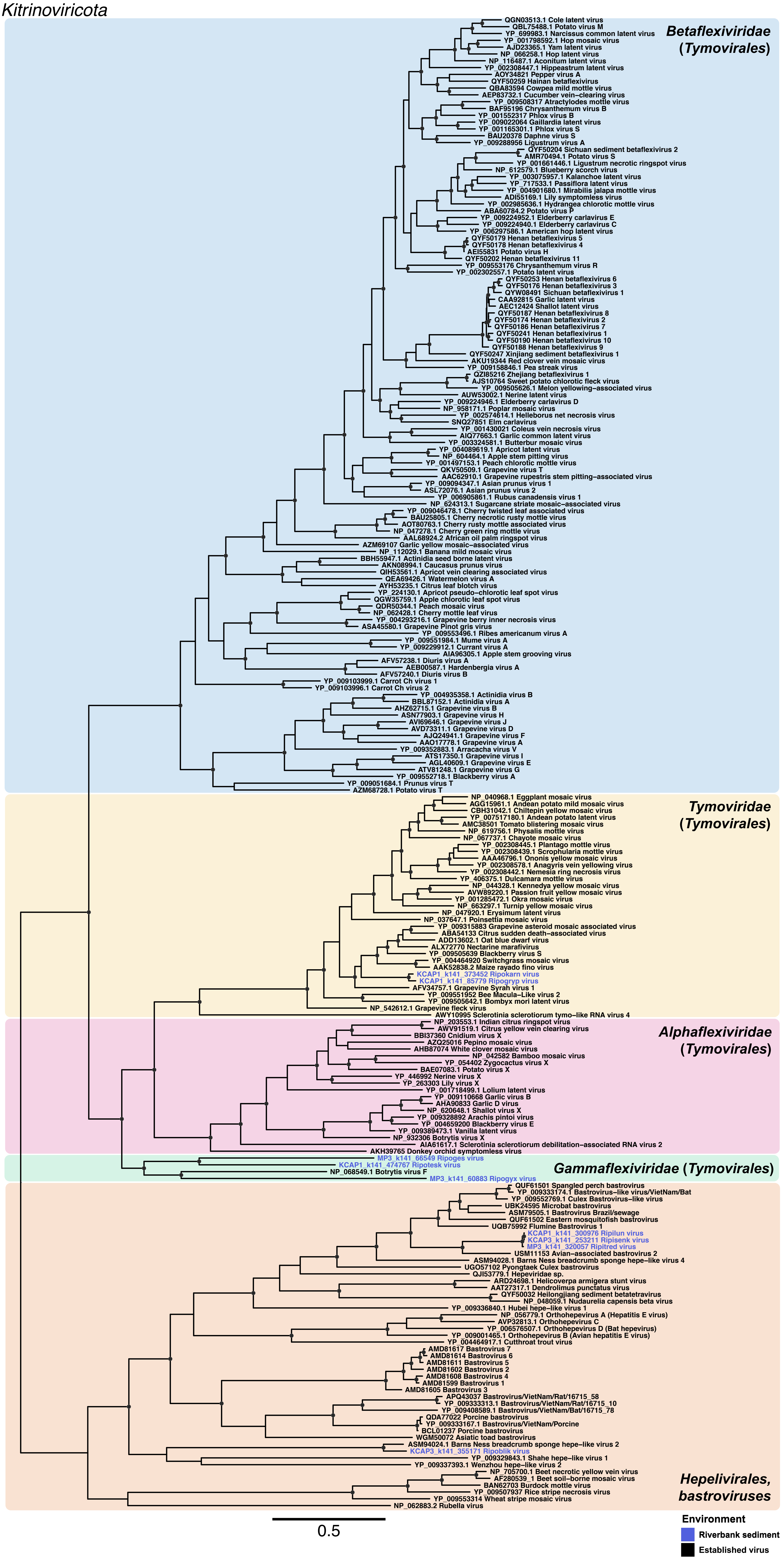

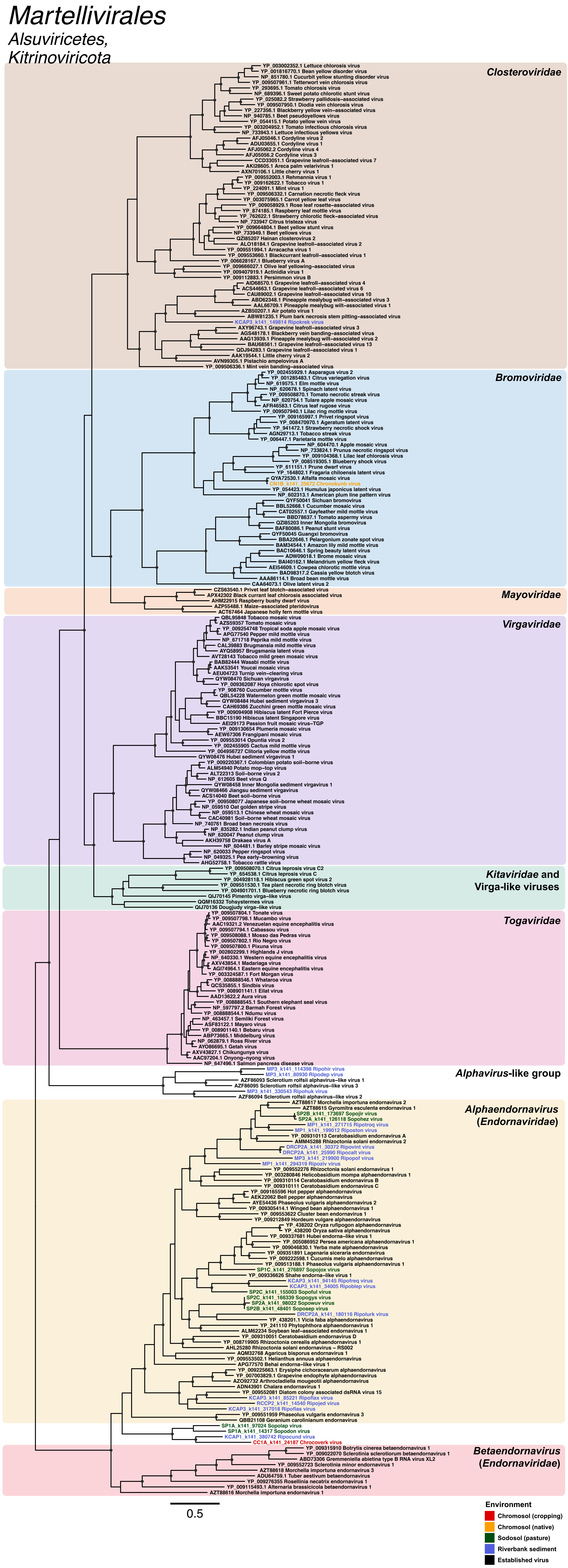

Martellivirales
Alsuviricetes,
Kitrinoviricota
Closteroviridae
YP_003002352.1 Lettuce chlorosis virus
YP_001816770.1 Bean yellow disorder virus
NP_851780.1 Cucurbit yellow stunting disorder virus
YP_009507961.1 Tetterwort vein chlorosis virus
YP_293695.1 Tomato chlorosis virus
NP_689396.1 Sweet potato chlorotic stunt virus
YP_025082.2 Strawberry pallidosis-associated virus
YP_009507950.1 Diodia vein chlorosis virus
YP_227356.1 Blackberry yellow vein-associated virus
NP_940785.1 Beet pseudoyellows virus
YP_054415.1 Potato yellow vein virus
YP_003204952.1 Tomato infectious chlorosis virus
NP_733943.1 Lettuce infectious yellows virus
AFJ05046.1 Cordyline virus 2
ADU03655.1 Cordyline virus 1
AFJ05062.2 Cordyline virus 4
AFJ05056.2 Cordyline virus 3
CCD33051.1 Grapevine leafroll-associated virus 7
AKI28605.1 Areca palm velarivirus 1
AXN70106.1 Little cherry virus 1
YP_009552003.1 Rehmannia virus 1
YP_009162622.1 Tobacco virus 1
YP_224091.1 Mint virus 1
YP_009506332.1 Carnation necrotic fleck virus
YP_003075965.1 Carrot yellow leaf virus
YP_009058929.1 Rose leaf rosette-associated virus
YP_874185.1 Raspberry leaf mottle virus
YP_762622.1 Strawberry chlorotic fleck-associated virus
NP_733947 Citrus tristeza virus
YP_009664804.1 Beet yellow stunt virus
NP_733949.1 Beet yellows virus
QZI85207 Hainan closterovirus 2
ALO18184.1 Grapevine leafroll-associated virus 2
YP_009551994.1 Arracacha virus 1
YP_009553660.1 Blackcurrant leafroll-associated virus 1
YP_006628167.1 Blueberry virus A
YP_009666027.1 Olive leaf yellowing-associated virus
YP_009407919.1 Actinidia virus 1
YP_009112883.1 Persimmon virus B
AID68570.1 Grapevine leafroll-associated virus 4
ACS44663.1 Grapevine leafroll-associated virus 6
CAU89002.1 Grapevine leafroll-associated virus 10
ABD62348.1 Pineapple mealybug wilt-associated virus 3
AAL66709.1 Pineapple mealybug wilt-associated virus 1
AZB50207.1 Air potato virus 1
ABW81235.1 Plum bark necrosis stem pitting-associated virus
KCAP3_k141_149814 Ripokrek virus
AXY96743.1 Grapevine leafroll-associated virus 3
AGS48178.1 Blackberry vein banding-associated virus
AAG13939.1 Pineapple mealybug wilt-associated virus 2
BAU68561.1 Grapevine leafroll-associated virus 13
QDJ94283.1 Grapevine leafroll-associated virus 1
AAK19544.1 Little cherry virus 2
AVN99305.1 Pistachio ampelovirus A
YP_009506336.1 Mint vein banding-associated virus
Bromoviridae
YP_002455929.1 Asparagus virus 2
YP_001285483.1 Citrus variegation virus
NP_619575.1 Elm mottle virus
NP_620678.1 Spinach latent virus
YP_009508870.1 Tomato necrotic streak virus
NP_620754.1 Tulare apple mosaic virus
AFR46583.1 Citrus leaf rugose virus
YP_009507940.1 Lilac ring mottle virus
YP_009165997.1 Privet ringspot virus
YP_008470970.1 Ageratum latent virus
YP_941472.1 Strawberry necrotic shock virus
AGN29713.1 Tobacco streak virus
YP_006447.1 Parietaria mottle virus
NP_604470.1 Apple mosaic virus
NP_733824.1 Prunus necrotic ringspot virus
YP_009104368.1 Lilac leaf chlorosis virus
YP_008519305.1 Blueberry shock virus
YP_611151.1 Prune dwarf virus
YP_164802.1 Fragaria chiloensis latent virus
QYA72530.1 Alfalfa mosaic virus
CN1B_k141_25672 Chronokunb virus
YP_054423.1 Humulus japonicus latent virus
NP_602313.1 American plum line pattern virus
QYF50041 Sichuan bromovirus
BBL52668.1 Cucumber mosaic virus
CAT02557.1 Gayfeather mild mottle virus
BBD78637.1 Tomato aspermy virus
QZI85203 Inner Mongolia bromovirus
BAF80086.1 Peanut stunt virus
QYF50045 Guangxi bromovirus
BBA22646.1 Pelargonium zonate spot virus
BAM34544.1 Amazon lily mild mottle virus
BAC10646.1 Spring beauty latent virus
ADW09018.1 Brome mosaic virus
BAI40162.1 Melandrium yellow fleck virus
AEI54609.1 Cowpea chlorotic mottle virus
BAD98317.2 Cassia yellow blotch virus
AAA86114.1 Broad bean mottle virus
CAA64073.1 Olive latent virus 2
Mayoviridae
CZS63540.1 Privet leaf blotch-associated virus
APX42302 Black currant leaf chlorosis associated virus
AHM22915 Raspberry bushy dwarf virus
AZP55488.1 Maize-associated pteridovirus
ACT67464 Japanese holly fern mottle virus
Virgaviridae
QBL95848 Tobacco mosaic virus
AZS59357 Tomato mosaic virus
YP_009254748 Tropical soda apple mosaic virus
APG77540 Pepper mild mottle virus
NP_671718 Paprika mild mottle virus
CAL39883 Brugmansia mild mottle virus
AYQ58957 Brugmansia latent virus
AVT28143 Tobacco mild green mosaic virus
BAB82444 Wasabi mottle virus
AAK53541 Youcai mosaic virus
AEU04723 Turnip vein-clearing virus
QYW08470 Sichuan virgavirus
YP_009362087 Hoya chlorotic spot virus
YP_908760 Cucumber mottle virus
QBL54228 Watermelon green mottle mosaic virus
QYW08484 Hubei sediment virgavirus 3
CAH69386 Zucchini green mottle mosaic virus
YP_009094908 Hibiscus latent Fort Pierce virus
BBC15190 Hibiscus latent Singapore virus
AEI29173 Passion fruit mosaic virus-TGP
YP_009130654 Plumeria mosaic virus
AEW67306 Frangipani mosaic virus
YP_009553014 Opuntia virus 2
YP_002455905 Cactus mild mottle virus
YP_004956727 Clitoria yellow mottle virus
QYW08476 Hubei sediment virgavirus 1
YP_009220367.1 Colombian potato soil-borne virus
ALM54940 Potato mop-top virus
ALT22313 Soil-borne virus 2
NP_612605 Beet virus Q
QYW08458 Inner Mongolia sediment virgavirus 1
QYW08466 Jiangsu sediment virgavirus
ACS14040 Beet soil-borne virus
YP_009508077 Japanese soil-borne wheat mosaic virus
NP_059510 Oat golden stripe virus
NP_059513.1 Chinese wheat mosaic virus
CAC40981 Soil-borne wheat mosaic virus
NP_740761 Broad bean necrosis virus
NP_835282.1 Indian peanut clump virus
NP_620047 Peanut clump virus
AKH39758 Drakaea virus A
NP_604481.1 Barley stripe mosaic virus
NP_620033 Pepper ringspot virus
NP_049325.1 Pea early-browning virus
AHG52758.1 Tobacco rattle virus
Kitaviridae and
Virga-like viruses
YP_009508070.1 Citrus leprosis virus C2
YP_654538.1 Citrus leprosis virus C
YP_004928118.1 Hibiscus green spot virus 2
YP_009551530.1 Tea plant necrotic ring blotch virus
YP_004901701.1 Blueberry necrotic ring blotch virus
QIJ70145 Pimento virga-like virus
QQM16332 Tohsystermes virus
QIJ70136 Dougjudy virga-like virus
Togaviridae
YP_009507804.1 Tonate virus
YP_009507798.1 Mucambo virus
AAC19321.2 Venezuelan equine encephalitis virus
YP_009507794.1 Cabassou virus
YP_009508088.1 Mosso das Pedras virus
YP_009507802.1 Rio Negro virus
YP_009507800.1 Pixuna virus
YP_002802299.1 Highlands J virus
NP_640330.1 Western equine encephalitis virus
AXV43854.1 Madariaga virus
AGI74964.1 Eastern equine encephalitis virus
YP_003324587.1 Fort Morgan virus
YP_008888546.1 Whataroa virus
QCS35855.1 Sindbis virus
YP_008901141.1 Eilat virus
AAD13622.2 Aura virus
YP_008888545.1 Southern elephant seal virus
NP_597797.2 Barmah Forest virus
YP_008888544.1 Ndumu virus
NP_463457.1 Semliki Forest virus
ASF83122.1 Mayaro virus
YP_008901140.1 Bebaru virus
ABP73665.1 Middelburg virus
NP_062879.1 Ross River virus
AYO86695.1 Getah virus
AXV43827.1 Chikungunya virus
AAC97204.1 Onyong-nyong virus
NP_647496.1 Salmon pancreas disease virus
Alphavirus-like group
MP3_k141_114398 Ripohir virus
MP3_k141_80930 Ripodue virus
AZF86093 Sclerotium rolfsii alphavirus-like virus 1
AZF86095 Sclerotium rolfsii alphavirus-like virus 3
MP3_k141_330543 Ripohuk virus
AZF86094 Sclerotium rolfsii alphavirus-like virus 2
Alphaendornavirus
(Endornaviridae)
AZT88617 Morchella importuna endornavirus 2
AZT88615 Gyromitra esculenta endornavirus 1
SP2B_k141_173697 Sopojir virus
SP2A_k141_126118 Sopohez virus
MP1_k141_271715 Ripotroq virus
MP1_k141_199012 Riposton virus
YP_009310113 Ceratobasidium endornavirus A
AMM45288 Rhizoctonia solani endornavirus 2
DRCP2A_k141_30372 Ripovint virus
DRCP2A_k141_25990 Ripocalt virus
MP3_k141_219900 Ripopof virus
MP1_k141_294319 Ripoziv virus
YP_009552276 Rhizoctonia solani endornavirus 1
YP_003280846 Helicobasidium mompa alphaendornavirus 1
YP_009310114 Ceratobasidium endornavirus B
YP_009310111 Ceratobasidium endornavirus C
YP_009165596 Hot pepper alphaendornavirus
AEK22062 Bell pepper alphaendornavirus
AYE54436 Phaseolus vulgaris alphaendornavirus 2
YP_009305414.1 Winged bean alphaendornavirus 1
YP_009553622 Cluster bean endornavirus 1
YP_009212849 Hordeum vulgare alphaendornavirus
YP_438202 Oryza rufipogon alphaendornavirus
YP_438200 Oryza sativa alphaendornavirus
YP_009337681 Hubei endorna-like virus 1
YP_005086952 Persea americana alphaendornavirus 1
YP_009046830.1 Yerba mate alphaendornavirus
YP_009351891 Lagenaria siceraria endornavirus
YP_009222598.1 Cucumis melo alphaendornavirus
YP_009513188.1 Phaseolus vulgaris alphaendornavirus 1
SP1C_k141_276897 Sopojox virus
YP_009336626 Shahe endorna-like virus 1
KCAP3_k141_94145 Ripofreq virus
KCAP3_k141_34005 Riposlep virus
SP2C_k141_155003 Sopoful virus
SP2C_k141_166339 Sopogys virus
SP2A_k141_98022 Sopowuv virus
SP2B_k141_48401 Soposep virus
DRCP2A_k141_180116 Ripolurk virus
YP_438201.1 Vicia faba alphaendornavirus
YP_241110 Phytophthora alphaendornavirus 1
ALM62234 Soybean leaf-associated endornavirus 1
YP_009310051 Ceratobasidium endornavirus D
YP_008719905 Rhizoctonia cerealis alphaendornavirus 1
AHL25280 Rhizoctonia solani endornavirus - RS002
AQM32768 Agaricus bisporus endornavirus 1
YP_009553502.1 Helianthus annuus alphaendornavirus
APG77570 Behai endorna-like virus 1
YP_009225663.1 Erysiphe cichoracearum alphaendornavirus
YP_007003829.1 Grapevine endophyte alphaendornavirus
AZO92732 Arthrocladiella mougeotii alphaendornavirus
ADN43901 Chalara endornavirus 1
YP_009552081 Diatom colony associated dsRNA virus 15
KCAP3_k141_85221 Ripoflax virus
RCCP2_k141_14540 Ripojed virus
KCAP3_k141_317018 Ripoflas virus
YP_009551959 Phaseolus vulgaris endornavirus 3
QBB21108 Geranium carolinianum endornavirus
SP1A_k141_97024 Sopolap virus
SP1A_k141_14317 Sopodon virus
KCAP1_k141_380742 Ripocund virus
CC1A_k141_24187 Chrocoverk virus
Betaendornavirus
(Endornaviridae)
YP_009315910 Botrytis cinerea betaendornavirus 1
YP_009022070 Sclerotinia sclerotiorum betaendornavirus 1
ABD73306 Gremmeniella abietina type B RNA virus XL2
YP_009552723 Sclerotinia minor endornavirus 1
AZT88618 Morchella importuna endornavirus 3
ADU64759.1 Tuber aestivum betaendornavirus
YP_009276355 Rosellinia necatrix endornavirus 1
YP_009115493.1 Alternaria brassicicola betaendornavirus 1
AZT88616 Morchella importuna endornavirus 1
0.5
Environment
Chromosol (cropping)
Chromosol (native)
Sodosol (pasture)
Riverbank sediment
Established virus

### *Nodamuvirales*

*Magsaviricetes, Kitrinoviricota*

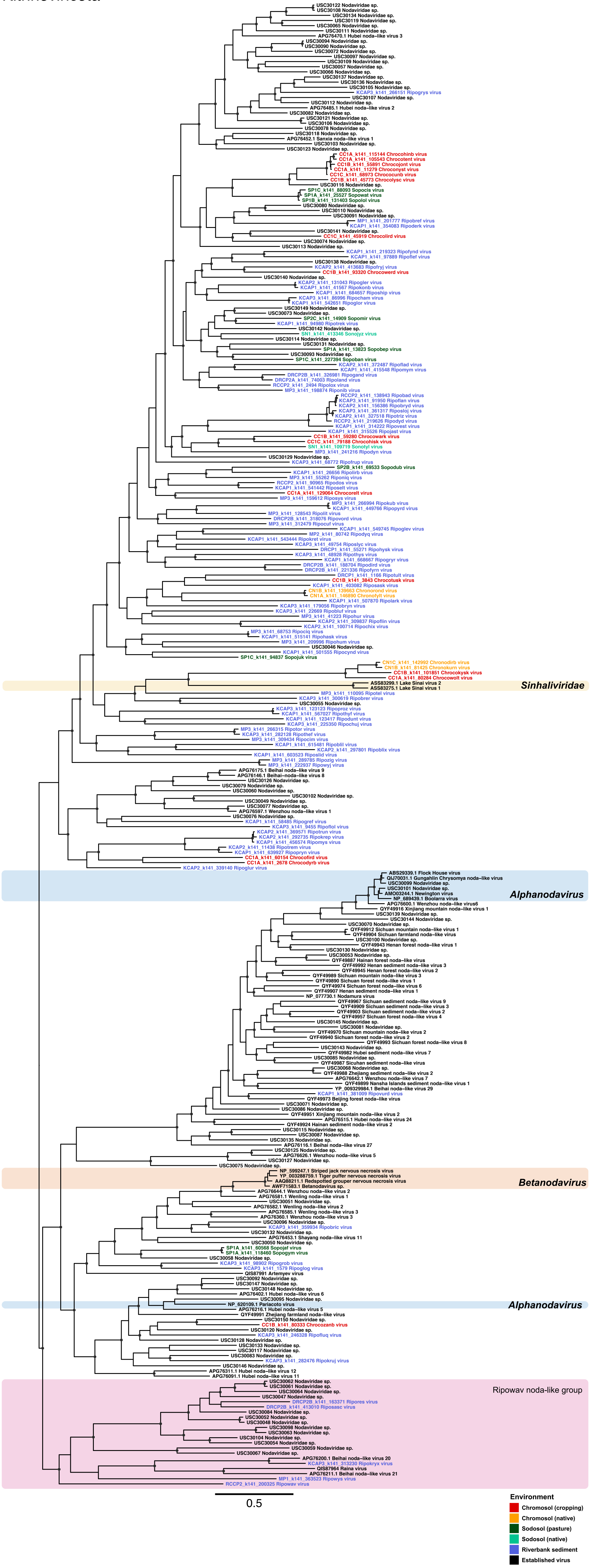

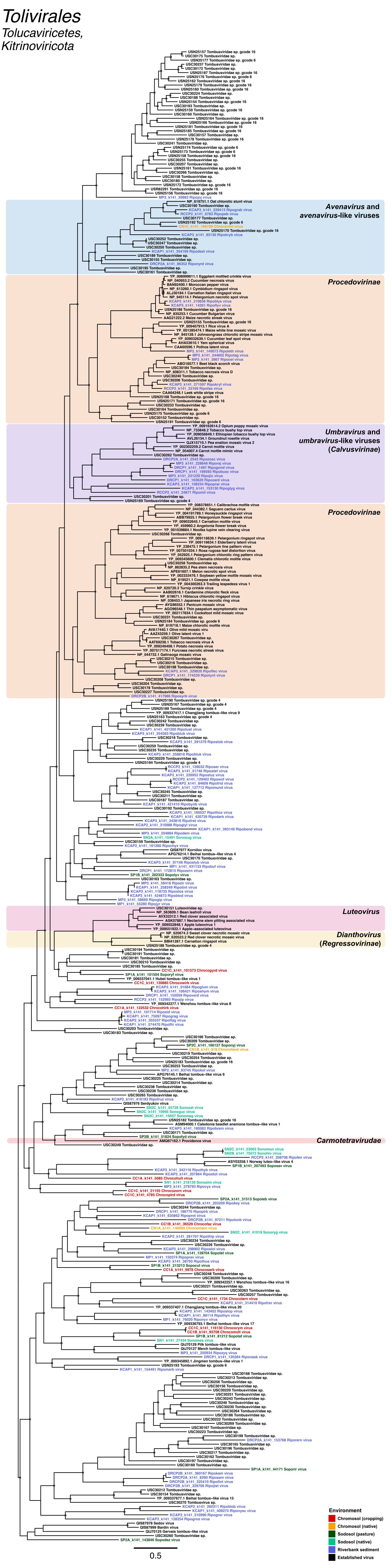

Tolivirales
Tolucaviricetes,
Kitrinoviricota
Avenavirus and avenavirus-like viruses
Procedovirinae
Umbravirus and umbravirus-like viruses (Calvusvirinae)
Procedovirinae
Luteovirus
Dianthovirus (Regressovirinae)
Carmotetraviridae
Environment
Chromosol (cropping)
Chromosol (native)
Sodosol (pasture)
Sodosol (native)
Riverbank sediment
Established virus
0.5

*Leviviricetes*

*Lenarviricota*

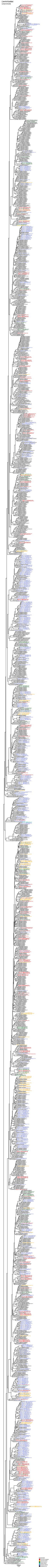

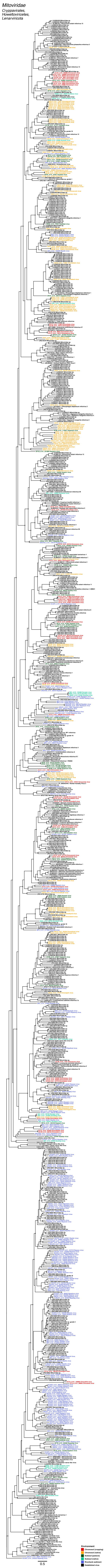
Mitoviridae
Cryppavirales,
Howeltoviricetes,
Lenarviricota
Environment
Chromosol (cropping)
Chromosol (native)
Sodosol (pasture)
Sodosol (native)
Riverbank sediment
Established virus
0.5

*Amabiliviricetes*

*Lenarviricota*

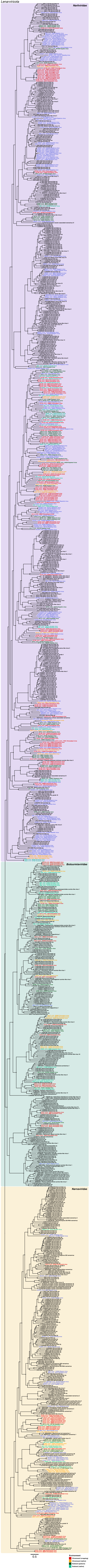

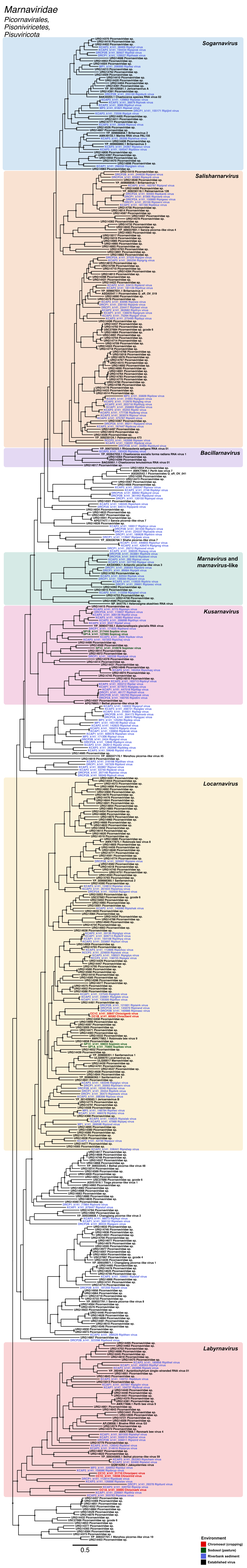

Marnaviridae
Picornavirales,
Pisoniviricetes,
Pisuviricota
Sogarnavirus
Salisharnavirus
Bacillarnavirus
Marnavirus and
marnavirus-like
Kusarnavirus
Locarnavirus
Labyrnavirus
0.5
Environment
Chromosol (cropping)
Sodosol (pasture)
Riverbank sediment
Established virus

### Picornaviridae, Polycipiviridae, Solinviviridae

Picornavirales, Pisoniviricetes, Pisuviricota

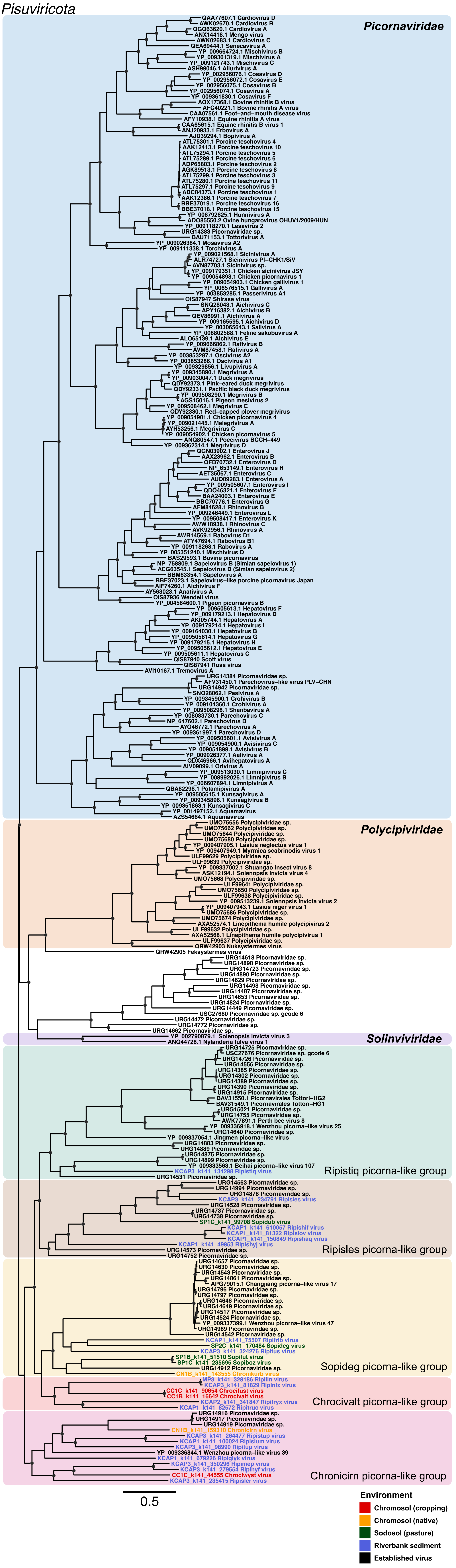

*Iflaviridae, Secoviridae, Dicistroviridae*

*Picornavirales, Pisoniviricetes, Pisuviricota*

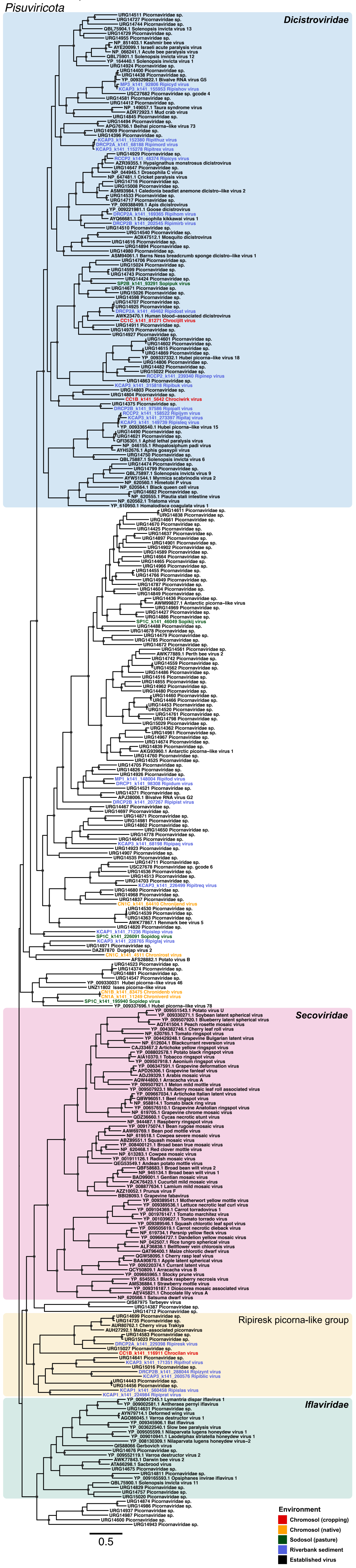

### *Sobelivirales*

*Pisoniviricetes,*
*Pisuviricota*

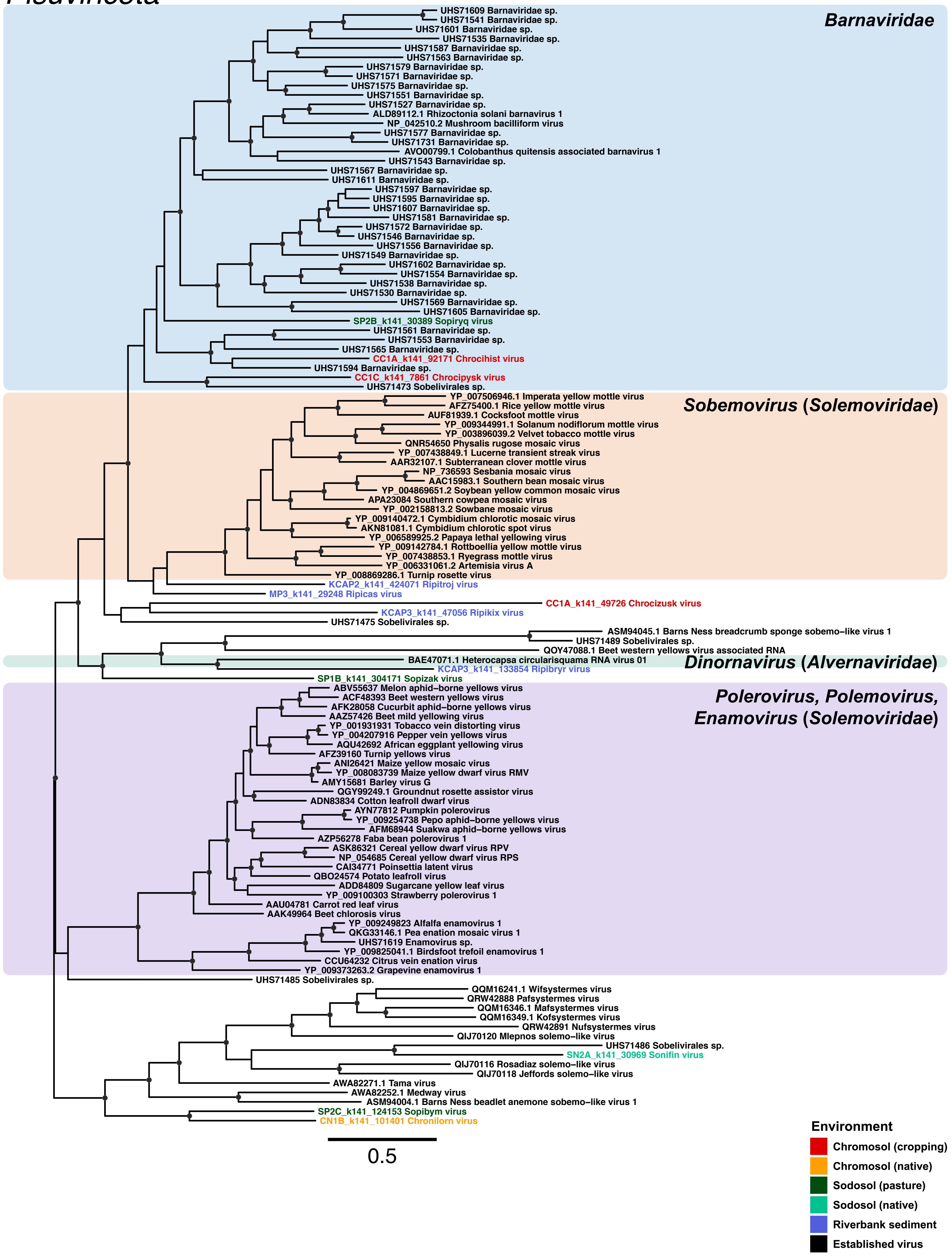

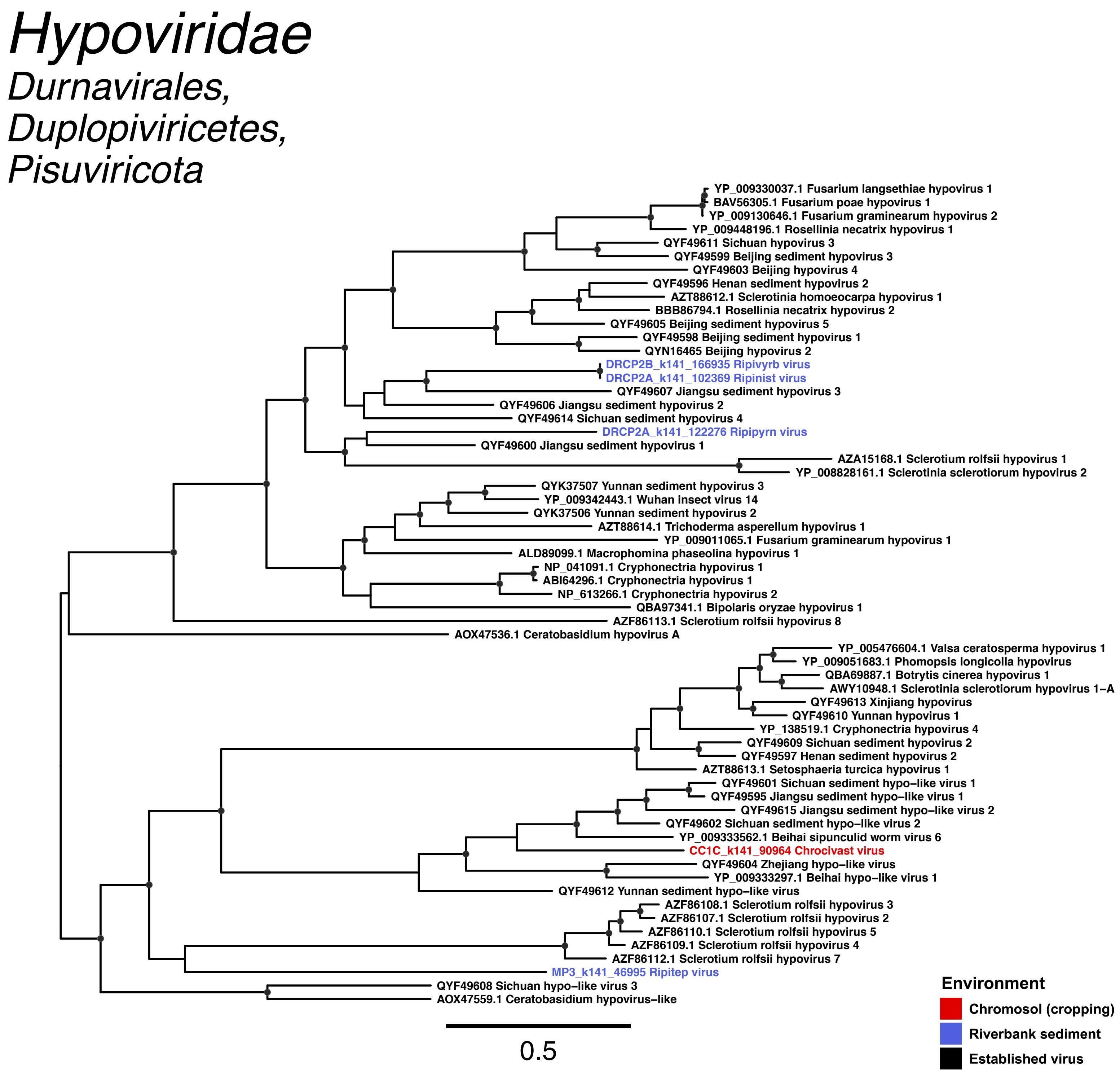
Hypoviridae
Durnavirales,
Duplopiviricetes,
Pisuviricota
YP_009330037.1 Fusarium langsethiae hypovirus 1
BAV56305.1 Fusarium poae hypovirus 1
YP_009130646.1 Fusarium graminearum hypovirus 2
YP_009448196.1 Rosellinia necatrix hypovirus 1
QYF49611 Sichuan hypovirus 3
QYF49599 Beijing sediment hypovirus 3
QYF49603 Beijing hypovirus 4
QYF49596 Henan sediment hypovirus 2
AZT88612.1 Sclerotinia homoeocarpa hypovirus 1
BBB86794.1 Rosellinia necatrix hypovirus 2
QYF49605 Beijing sediment hypovirus 5
QYF49598 Beijing sediment hypovirus 1
QYN16465 Beijing hypovirus 2
DRCP2B_k141_166935 Ripivyrb virus
DRCP2A_k141_102369 Ripinist virus
QYF49607 Jiangsu sediment hypovirus 3
QYF49606 Jiangsu sediment hypovirus 2
QYF49614 Sichuan sediment hypovirus 4
DRCP2A_k141_122276 Ripipyrn virus
QYF49600 Jiangsu sediment hypovirus 1
AZA15168.1 Sclerotium rolfsii hypovirus 1
YP_008828161.1 Sclerotinia sclerotiorum hypovirus 2
QYK37507 Yunnan sediment hypovirus 3
YP_009342443.1 Wuhan insect virus 14
QYK37506 Yunnan sediment hypovirus 2
AZT88614.1 Trichoderma asperellum hypovirus 1
YP_009011065.1 Fusarium graminearum hypovirus 1
ALD89099.1 Macrophomina phaseolina hypovirus 1
NP_041091.1 Cryphonectria hypovirus 1
ABI64296.1 Cryphonectria hypovirus 1
NP_613266.1 Cryphonectria hypovirus 2
QBA97341.1 Bipolaris oryzae hypovirus 1
AZF86113.1 Sclerotium rolfsii hypovirus 8
AOX47536.1 Ceratobasidium hypovirus A
YP_005476604.1 Valsa ceratosperma hypovirus 1
YP_009051683.1 Phomopsis longicolla hypovirus
QBA69887.1 Botrytis cinerea hypovirus 1
AWY10948.1 Sclerotinia sclerotiorum hypovirus 1–A
QYF49613 Xinjiang hypovirus
QYF49610 Yunnan hypovirus 1
YP_138519.1 Cryphonectria hypovirus 4
QYF49609 Sichuan sediment hypovirus 2
QYF49597 Henan sediment hypovirus 2
AZT88613.1 Setosphaeria turcica hypovirus 1
QYF49601 Sichuan sediment hypo–like virus 1
QYF49595 Jiangsu sediment hypo–like virus 1
QYF49615 Jiangsu sediment hypo–like virus 2
QYF49602 Sichuan sediment hypo–like virus 2
YP_009333562.1 Beihai sipunculid worm virus 6
CC1C_k141_90964 Chrocivast virus
QYF49604 Zhejiang hypo–like virus
YP_009333297.1 Beihai hypo–like virus 1
QYF49612 Yunnan sediment hypo–like virus
AZF86108.1 Sclerotium rolfsii hypovirus 3
AZF86107.1 Sclerotium rolfsii hypovirus 2
AZF86110.1 Sclerotium rolfsii hypovirus 5
AZF86109.1 Sclerotium rolfsii hypovirus 4
AZF86112.1 Sclerotium rolfsii hypovirus 7
MP3_k141_46995 Ripitep virus
QYF49608 Sichuan hypo–like virus 3
AOX47559.1 Ceratobasidium hypovirus–like
0.5
Environment
Chromosol (cropping)
Riverbank sediment
Established virus

### Partiti–Picobirna

*Durnavirales,*
*Duplopiviricetes,*
*Pisuviricota*

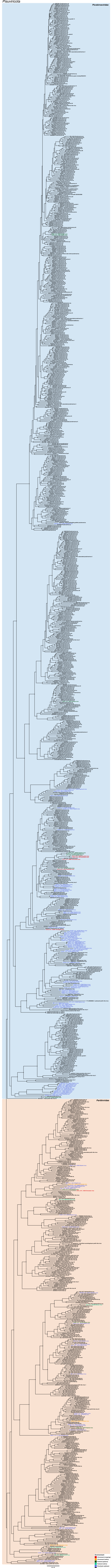

### *Monjiviricetes*

#### *Negarnaviricota*

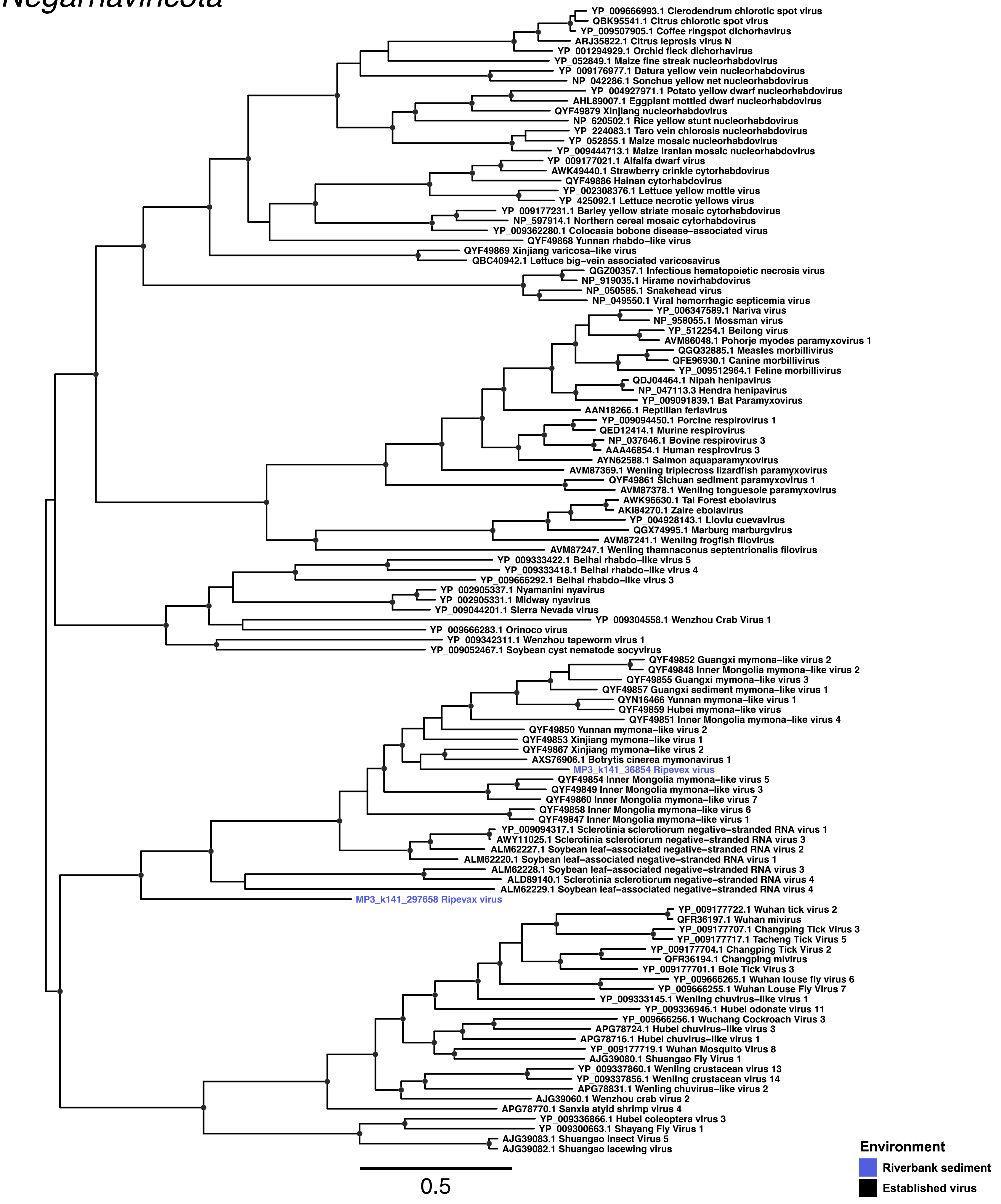

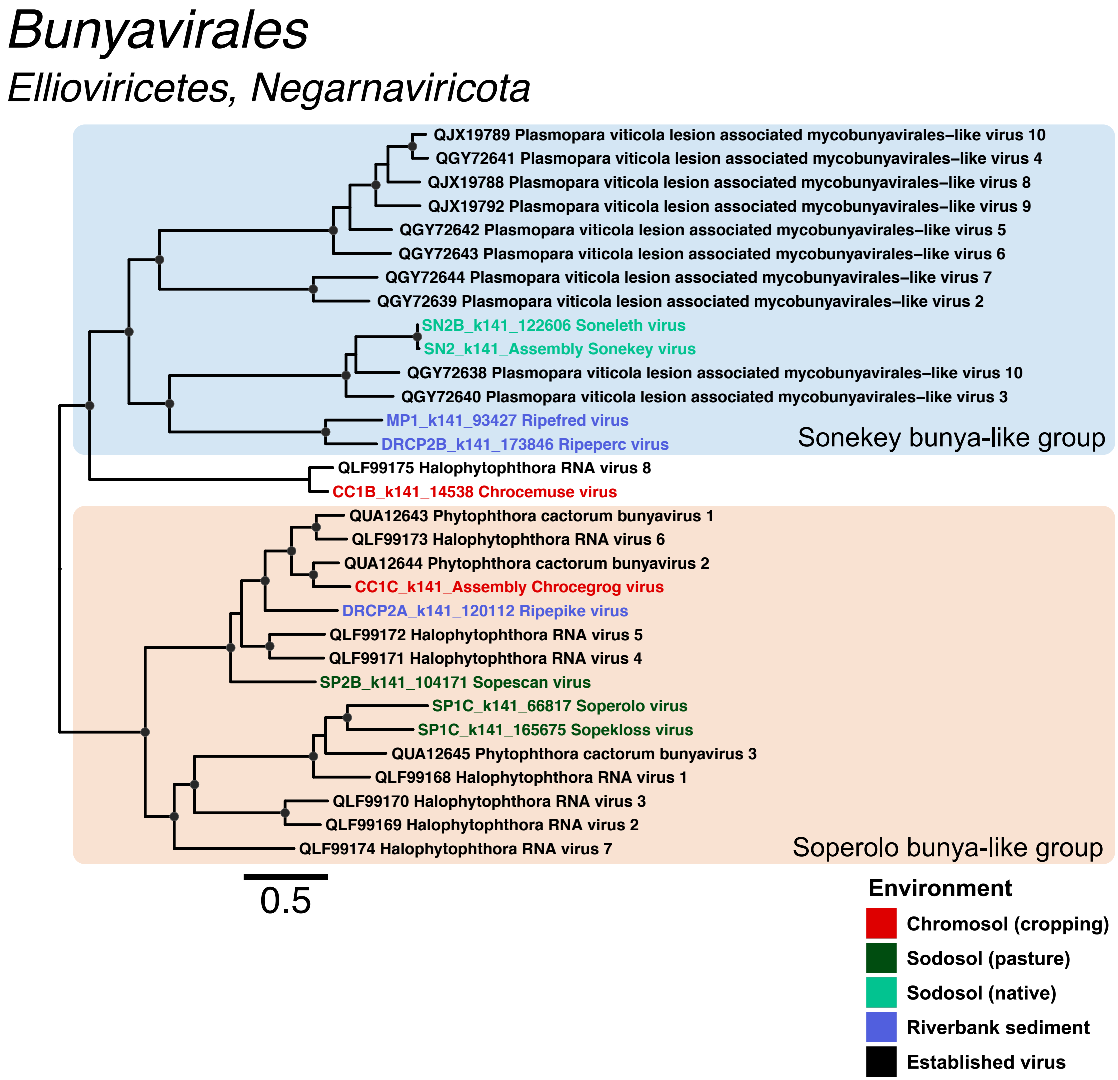
Bunyavirales
Elliovirícetes, Negarnaviricota
QJX19789 Plasmopara viticola lesion associated mycobunyavirales–like virus 10
QGY72641 Plasmopara viticola lesion associated mycobunyavirales–like virus 4
QJX19788 Plasmopara viticola lesion associated mycobunyavirales–like virus 8
QJX19792 Plasmopara viticola lesion associated mycobunyavirales–like virus 9
QGY72642 Plasmopara viticola lesion associated mycobunyavirales–like virus 5
QGY72643 Plasmopara viticola lesion associated mycobunyavirales–like virus 6
QGY72644 Plasmopara viticola lesion associated mycobunyavirales–like virus 7
QGY72639 Plasmopara viticola lesion associated mycobunyavirales–like virus 2
SN2B_k141_122606 Soneleth virus
SN2_k141_Assembly Sonekey virus
QGY72638 Plasmopara viticola lesion associated mycobunyavirales–like virus 10
QGY72640 Plasmopara viticola lesion associated mycobunyavirales–like virus 3
MP1_k141_93427 Ripefred virus
DRCP2B_k141_173846 Ripeperc virus
Sonekey bunya-like group
QLF99175 Halophytophthora RNA virus 8
CC1B_k141_14538 Chrocemuse virus
QUA12643 Phytophthora cactorum bunyavirus 1
QLF99173 Halophytophthora RNA virus 6
QUA12644 Phytophthora cactorum bunyavirus 2
CC1C_k141_Assembly Chrocegrog virus
DRCP2A_k141_120112 Ripepike virus
QLF99172 Halophytophthora RNA virus 5
QLF99171 Halophytophthora RNA virus 4
SP2B_k141_104171 Sopescan virus
SP1C_k141_66817 Soperolo virus
SP1C_k141_165675 Sopekloss virus
QUA12645 Phytophthora cactorum bunyavirus 3
QLF99168 Halophytophthora RNA virus 1
QLF99170 Halophytophthora RNA virus 3
QLF99169 Halophytophthora RNA virus 2
QLF99174 Halophytophthora RNA virus 7
Soperolo bunya-like group
0.5
Environment
Chromosol (cropping)
Sodosol (pasture)
Sodosol (native)
Riverbank sediment
Established virus

### *Cystoviridae*

*Mindivirales, Vidaverviricetes, Duplornaviricota*

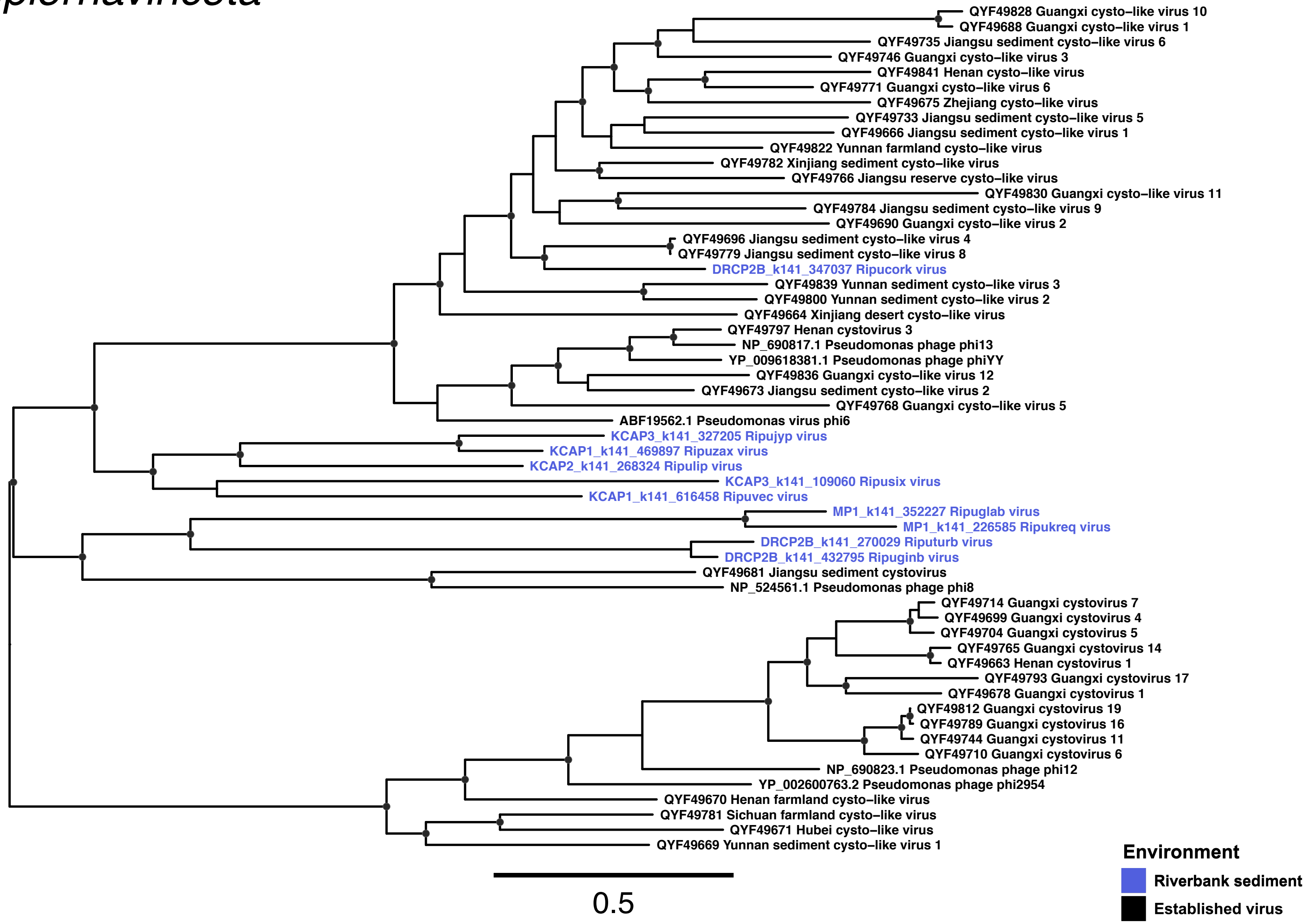

### *Reovirales*

#### *Resentoviricetes, Duplornaviricota*

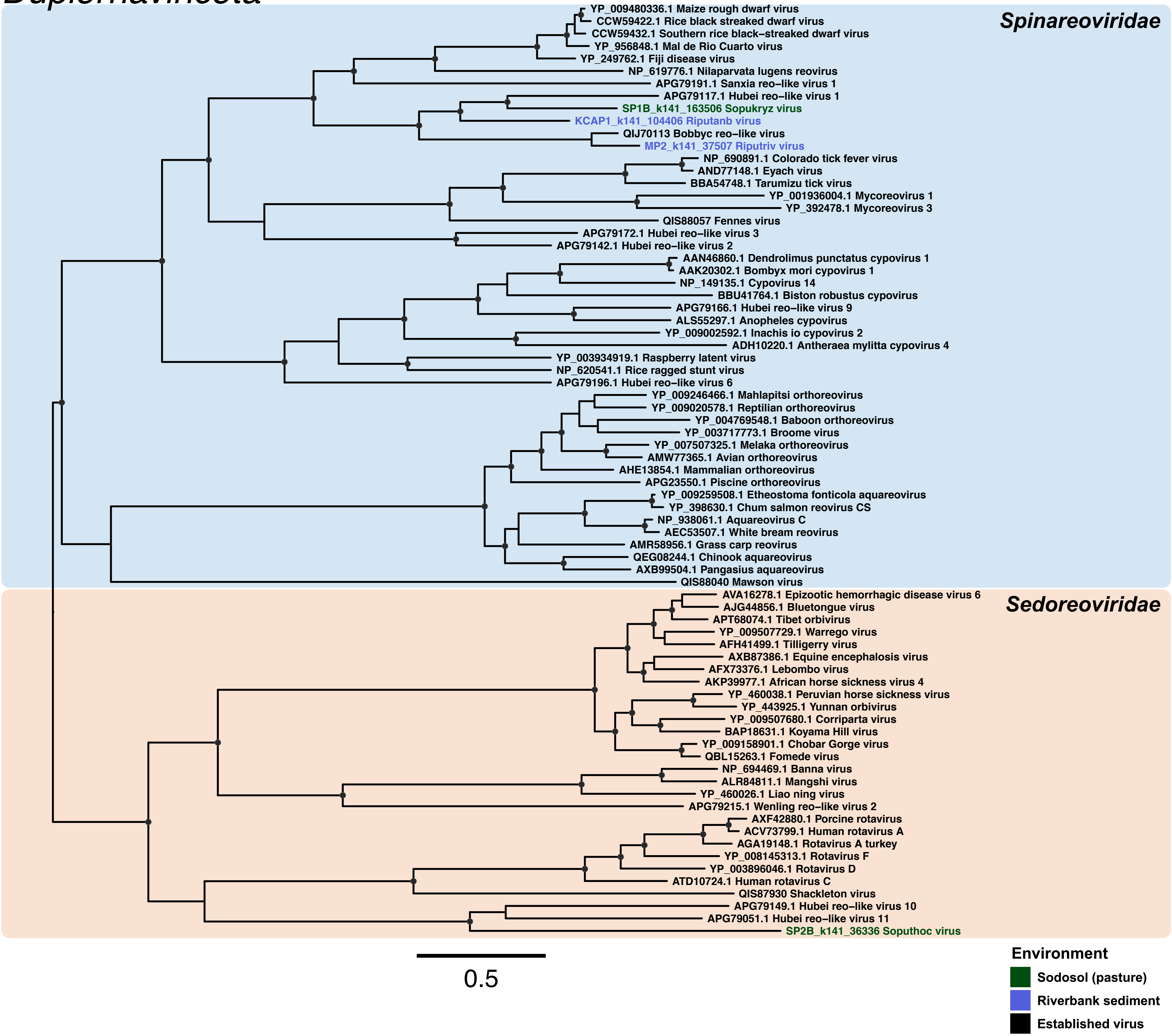

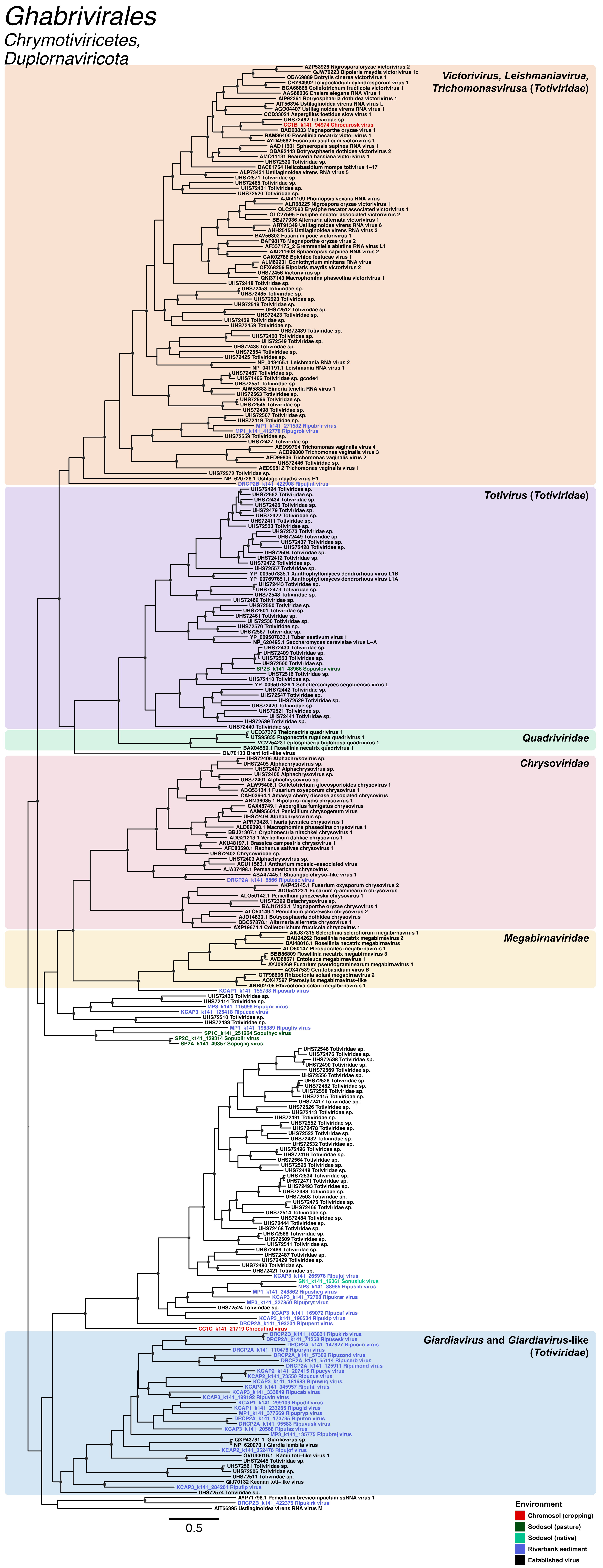
Ghabrivirales
Chrymotiviricetes,
Duplornaviricota
Victorivirus, Leishmaniavirua, Trichomonasvirusa (Totiviridae)
AZP53926 Nigrospora oryzae victorivirus 2
QJW70223 Bipolaris maydis victorivirus 1c
QBA69889 Botrytis cinerea victorivirus 1
CBY84992 Tolypocladium cylindrosporum virus 1
BCA66668 Colletotrichum fructicola victorivirus 1
AAS68036 Chalara elegans RNA Virus 1
AIP92361 Botryosphaeria dothidea victorivirus 1
AIT56394 Ustilaginoidea virens RNA virus L
AGO04407 Ustilaginoidea virens RNA virus 1
CCD33024 Aspergillus foetidus slow virus 1
UHS72462 Totiviridae sp.
CC1B_k141_94974 Chrocurosk virus
BAD60833 Magnaporthe oryzae virus 1
BAM36400 Rosellinia necatrix victorivirus 1
AYD49682 Fusarium asiaticum victorivirus 1
AAD11601 Sphaeropsis sapinea RNA virus 1
QBA82443 Botryosphaeria dothidea victorivirus 2
AMQ11131 Beauveria bassiana victorivirus 1
UHS72530 Totiviridae sp.
BAC81754 Helicobasidium mompa totivirus 1–17
ALP73431 Ustilaginoidea virens RNA virus 5
UHS72571 Totiviridae sp.
UHS72465 Totiviridae sp.
UHS72431 Totiviridae sp.
UHS72520 Totiviridae sp.
AJA41109 Phomopsis vexans RNA virus
ALR68225 Nigrospora oryzae victorivirus 1
QLC27593 Erysiphe necator associated victorivirus 1
QLC27595 Erysiphe necator associated victorivirus 2
BBJ77936 Alternaria alternata victorivirus 1
ART91349 Ustilaginoidea virens RNA virus 6
AHH25155 Ustilaginoidea virens RNA virus 3
BAV56302 Fusarium poae victorivirus 1
BAF98178 Magnaporthe oryzae virus 2
AF337175_2 Gremmeniella abietina RNA virus L1
AAD11603 Sphaeropsis sapinea RNA virus 2
CAK02788 Epichloe festucae virus 1
ALM62231 Coniothyrium minitans RNA virus
QFX68259 Bipolaris maydis victorivirus 2
UHS72456 Victorivirus sp.
QKI37143 Macrophomina phaseolina victorivirus 1
UHS72418 Totiviridae sp.
UHS72453 Totiviridae sp.
UHS72485 Totiviridae sp.
UHS72523 Totiviridae sp.
UHS72519 Totiviridae sp.
UHS72512 Totiviridae sp.
UHS72423 Totiviridae sp.
UHS72439 Totiviridae sp.
UHS72459 Totiviridae sp.
UHS72489 Totiviridae sp.
UHS72460 Totiviridae sp.
UHS72549 Totiviridae sp.
UHS72438 Totiviridae sp.
UHS72554 Totiviridae sp.
UHS72425 Totiviridae sp.
NP_043465.1 Leishmania RNA virus 2
NP_041191.1 Leishmania RNA virus 1
UHS72467 Totiviridae sp.
UHS71466 Totiviridae sp. gcode4
UHS72551 Totiviridae sp.
AIW58883 Eimeria tenella RNA virus 1
UHS72563 Totiviridae sp.
UHS72566 Totiviridae sp.
UHS72545 Totiviridae sp.
UHS72498 Totiviridae sp.
UHS72507 Totiviridae sp.
UHS72419 Totiviridae sp.
MP1_k141_271532 Ripubrir virus
MP1_k141_412778 Ripugrok virus
UHS72559 Totiviridae sp.
UHS72427 Totiviridae sp.
AED99794 Trichomonas vaginalis virus 4
AED99800 Trichomonas vaginalis virus 3
AED99806 Trichomonas vaginalis virus 2
UHS72446 Totiviridae sp.
AED99812 Trichomonas vaginalis virus 1
UHS72572 Totiviridae sp.
NP_620728.1 Ustilago maydis virus H1
DRCP2B_k141_422908 Ripujint virus
Totivirus (Totiviridae)
UHS72424 Totiviridae sp.
UHS72562 Totiviridae sp.
UHS72434 Totiviridae sp.
UHS72426 Totiviridae sp.
UHS72479 Totiviridae sp.
UHS72422 Totiviridae sp.
UHS72411 Totiviridae sp.
UHS72533 Totiviridae sp.
UHS72573 Totiviridae sp.
UHS72449 Totiviridae sp.
UHS72437 Totiviridae sp.
UHS72428 Totiviridae sp.
UHS72504 Totiviridae sp.
UHS72412 Totiviridae sp.
UHS72472 Totiviridae sp.
UHS72557 Totiviridae sp.
YP_009507835.1 Xanthophyllomyces dendrorhous virus L1B
YP_007697651.1 Xanthophyllomyces dendrorhous virus L1A
UHS72443 Totiviridae sp.
UHS72473 Totiviridae sp.
UHS72548 Totiviridae sp.
UHS72469 Totiviridae sp.
UHS72550 Totiviridae sp.
UHS72501 Totiviridae sp.
UHS72461 Totiviridae sp.
UHS72536 Totiviridae sp.
UHS72570 Totiviridae sp.
UHS72567 Totiviridae sp.
YP_009507833.1 Tuber aestivum virus 1
NP_620495.1 Saccharomyces cerevisiae virus L–A
UHS72430 Totiviridae sp.
UHS72409 Totiviridae sp.
UHS72553 Totiviridae sp.
UHS72500 Totiviridae sp.
SP2B_k141_48966 Sopuslov virus
UHS72516 Totiviridae sp.
UHS72410 Totiviridae sp.
YP_009507829.1 Scheffersomyces segobiensis virus L
UHS72442 Totiviridae sp.
UHS72547 Totiviridae sp.
UHS72529 Totiviridae sp.
UHS72420 Totiviridae sp.
UHS72521 Totiviridae sp.
UHS72441 Totiviridae sp.
UHS72539 Totiviridae sp.
UHS72440 Totiviridae sp.
Quadriviridae
UED37376 Thelonectria quadrivirus 1
UTS95835 Rugonectria rugulosa quadrivirus 1
VCV25423 Leptosphaeria biglobosa quadrivirus 1
BAX04559.1 Rosellinia necatrix quadrivirus 1
QIJ70133 Brent toti–like virus
Chrysoviridae
UHS72406 Alphachrysovirus sp.
UHS72405 Alphachrysovirus sp.
UHS72407 Alphachrysovirus sp.
UHS72400 Alphachrysovirus sp.
UHS72401 Alphachrysovirus sp.
ALW95408.1 Colletotrichum gloeosporioides chrysovirus 1
ABO53134.1 Fusarium oxysporum chrysovirus 1
CAH03664.1 Amasya cherry disease associated chrysovirus
ARM36035.1 Bipolaris maydis chrysovirus 1
CAX48749.1 Aspergillus fumigatus chrysovirus
AAM95601.1 Penicillium chrysogenum virus
UHS72404 Alphachrysovirus sp.
APR73428.1 Isaria javanica chrysovirus 1
ALD89090.1 Macrophomina phaseolina chrysovirus 1
BBJ21307.1 Cryphonectria nitschkei chrysovirus 1
ADG21213.1 Verticillium dahliae chrysovirus 1
AKU48197.1 Brassica campestris chrysovirus 1
AFE83590.1 Raphanus sativas chrysovirus 1
UHS72402 Chrysoviridae sp.
UHS72403 Alphachrysovirus sp.
ACU11563.1 Anthurium mosaic–associated virus
AJA37498.1 Persea americana chrysovirus
ASA47445.1 Shuangao chryso–like virus 1
DRCP2A_k141_6866 Riputesc virus
AKP45145.1 Fusarium oxysporum chrysovirus 2
ADU54123.1 Fusarium graminearum chrysovirus
ALO50142.1 Penicillium janczewskii chrysovirus 1
UHS72399 Betachrysovirus sp.
BAJ15133.1 Magnaporthe oryzae chrysovirus 1
ALO50149.1 Penicillium janczewskii chrysovirus 2
AJD14830.1 Botryosphaeria dothidea chrysovirus 1
BBC27878.1 Alternaria alternata chrysovirus 1
AXP19674.1 Colletotrichum fructicola chrysovirus 1
Megabirnaviridae
AKJ87315 Sclerotinia sclerotiorum megabirnavirus 1
BAU24262 Rosellinia necatrix megabirnavirus 2
BAI48016.1 Rosellinia necatrix megabirnavirus
ALO50147 Pleosporales megabirnavirus 1
BBB86809 Rosellinia necatrix megabirnavirus 3
AVD68671 Entoleuca megabirnavirus 1
AYJ09269 Fusarium pseudograminearum megabirnavirus 1
AOX47539 Ceratobasidium virus B
QTF98696 Rhizoctonia solani megabirnavirus 2
AOX47597 Pterostylis megabirnavirus–like
ANR02705 Rhizoctonia solani megabirnavirus 1
KCAP1_k141_155733 Ripusarb virus
UHS72436 Totiviridae sp.
UHS72414 Totiviridae sp.
MP3_k141_115098 Ripugrir virus
KCAP3_k141_125418 Ripucex virus
UHS72510 Totiviridae sp.
UHS72433 Totiviridae sp.
MP1_k141_198389 Ripuglis virus
SP1C_k141_251264 Soputhyc virus
SP2C_k141_129314 Sopubliir virus
SP2A_k141_49857 Sopuglig virus
UHS72546 Totiviridae sp.
UHS72476 Totiviridae sp.
UHS72538 Totiviridae sp.
UHS72490 Totiviridae sp.
UHS72569 Totiviridae sp.
UHS72556 Totiviridae sp.
UHS72528 Totiviridae sp.
UHS72482 Totiviridae sp.
UHS72558 Totiviridae sp.
UHS72415 Totiviridae sp.
UHS72417 Totiviridae sp.
UHS72526 Totiviridae sp.
UHS72413 Totiviridae sp.
UHS72491 Totiviridae sp.
UHS72552 Totiviridae sp.
UHS72478 Totiviridae sp.
UHS72522 Totiviridae sp.
UHS72432 Totiviridae sp.
UHS72532 Totiviridae sp.
UHS72496 Totiviridae sp.
UHS72416 Totiviridae sp.
UHS72564 Totiviridae sp.
UHS72525 Totiviridae sp.
UHS72448 Totiviridae sp.
UHS72534 Totiviridae sp.
UHS72471 Totiviridae sp.
UHS72493 Totiviridae sp.
UHS72483 Totiviridae sp.
UHS72503 Totiviridae sp.
UHS72475 Totiviridae sp.
UHS72466 Totiviridae sp.
UHS72514 Totiviridae sp.
UHS72484 Totiviridae sp.
UHS72444 Totiviridae sp.
UHS72468 Totiviridae sp.
UHS72568 Totiviridae sp.
UHS72509 Totiviridae sp.
UHS72541 Totiviridae sp.
UHS72488 Totiviridae sp.
UHS72487 Totiviridae sp.
UHS72429 Totiviridae sp.
UHS72480 Totiviridae sp.
UHS72421 Totiviridae sp.
KCAP3_k141_265976 Ripujoj virus
SN1_k141_16361 Sonusluk virus
MP3_k141_88965 Ripuslib virus
MP1_k141_348862 Ripusheg virus
KCAP3_k141_72708 Ripukrar virus
MP3_k141_327850 Ripupryt virus
UHS72524 Totiviridae sp.
KCAP3_k141_169072 Ripucaf virus
KCAP3_k141_196534 Ripukip virus
DRCP2A_k141_193204 Ripupent virus
CC1C_k141_21719 Chrocutind virus
Giardiavirus and Giardiavirus-like (Totiviridae)
DRCP2B_k141_103831 Ripukirb virus
DRCP2A_k141_71258 Ripusesk virus
DRCP2A_k141_147827 Ripucim virus
DRCP2A_k141_110478 Ripurym virus
DRCP2A_k141_57302 Ripuzond virus
DRCP2A_k141_55114 Ripucerb virus
DRCP2A_k141_125911 Ripumond virus
KCAP2_k141_207415 Ripucyv virus
KCAP2_k141_73550 Ripucus virus
KCAP3_k141_181683 Ripuwuq virus
KCAP3_k141_345957 Ripuhil virus
KCAP3_k141_333849 Ripucab virus
KCAP3_k141_199192 Ripuvin virus
KCAP1_k141_299109 Ripudil virus
KCAP1_k141_233265 Ripugid virus
MP1_k141_377669 Ripupryp virus
DRCP2A_k141_173735 Riputon virus
DRCP2A_k141_95583 Ripuvusk virus
KCAP3_k141_20568 Riputaz virus
MP3_k141_135775 Ripubrej virus
QXP43781.1 Giardiavirus sp.
NP_620070.1 Giardia lamblia virus
KCAP2_k141_352476 Ripujof virus
QVU40016.1 Kamu toti–like virus 1
UHS72445 Totiviridae sp.
UHS72561 Totiviridae sp.
UHS72506 Totiviridae sp.
UHS72511 Totiviridae sp.
QIJ70132 Keenan toti–like virus
KCAP3_k141_284261 Ripuflip virus
UHS72574 Totiviridae sp.
AYP71798.1 Penicillium brevicompactum ssRNA virus 1
DRCP2B_k141_422375 Ripukirk virus
AIT56395 Ustilaginoidea virens RNA virus M
0.5
Environment
Chromosol (cropping)
Sodosol (pasture)
Sodosol (native)
Riverbank sediment
Established virus
